# Stabilization of memory on neural manifolds through multiple synaptic time scales

**DOI:** 10.1101/2025.01.30.635710

**Authors:** Georg Chechelnizki, Nimrod Shaham, Alon Salhov, Yoram Burak

**Affiliations:** Edmond and Lily Safra Center for Brain Sciences, Jerusalem, Israel; Racah Institute of Physics, Jerusalem, Israel

## Abstract

Working memory is commonly thought to be implemented in the brain by the persistent activity of attractor neural networks. Compelling evidence exists for the implementation of such memory networks in vertebrate and invertebrate brains. Naively, retention of *continuous* memories in attractor networks – as observed in the head direction systems of mammals and insects and the oculomotor integrator in vertebrates – is particularly sensitive to neural stochasticity, because noise can continuously shift the activity of the network along its manifold of steady states. As these random drifts accumulate over time, the stored memory degrades. Small networks, such as the head direction system of insects, are particularly prone to this form of memory deterioration, and it is unclear how such networks can maintain their persistent state over behaviorally relevant timescales. Here we identify a neural mechanism that effectively counteracts random drifts by employing a combination of slow excitatory and fast inhibitory synaptic connections. We show that the proposed mechanism can substantially decrease diffusivity in linear and nonlinear networks. Hence, the mechanism offers a plausible explanation for how the brain can stably store memories of continuous quantities, despite the ubiquity of noise. Finally, we identify how to engineer connectivity such that variability in neural activity perpendicular to the attractor is unaffected by the stabilization mechanism. This scheme allows for tight confinement of neural activity patterns to a low dimensional manifold, as well as the rapid relaxation of transient modes of activity back into the attractor. The theory suggests that neurons in head direction cell networks that are commonly thought to be utilized for velocity integration may also aid in stabilization against noise-driven motion.

## Introduction

Living organisms rely on short-term memory for their survival – in navigation [1–5], decision making [6–9], sensorimotor integration [10, 11], and many other cognitive tasks. This highlights the importance of elucidating the mechanisms through which the brain represents information about the past in its ongoing dynamics. Continuous attractor neural networks (CANs) are an important class of recurrent neural network architectures that can maintain short-term memory of continuous variables. While such networks may comprise many neurons, their population activity is constrained to a low-dimensional manifold, such as a line [10–13], a ring [14–19], a plane [20–22] or a torus [23–25]. Each point on the attractor corresponds to a unique combination of firing rates that the network can sustain persistently, and each such state can be mapped to represent a particular value of the stored memory. Several neural circuits in living organisms are thought to be organized in CANs, among them: the oculomotor integrator [10–13], head direction cells in rodents [15, 26–29] and insects [16–18, 30–34] and grid cells in the entorhinal cortex [23, 24, 35–39]. Head direction cell systems, which appear to have independently emerged across diverse species, will serve as a prime example in this study.

While most of the theoretical studies on CANs modeled their activity in terms of deterministic neural dynamics, it has become clear in recent years that CANs are highly prone to neural noise [40, 41], an inherent aspect of neural dynamics in the brain [42], because memories can be continuously shifted by noise along the attractor without resistance. Several recent works provide empirical evidence for such random motion in specific attractor networks [15, 18, 43]. Termed *diffusion*, this phenomenon implies that memories within such systems should degrade over time.

Noise-driven diffusion is particularly pronounced in theoretical models when the number of neurons is small [40], yet there are examples of CANs in the brain that contain only tens or hundreds of neurons, e.g., the oculomotor system of fish [11] and the head direction system of the fly [44], where we demonstrate below that noise is naively expected to drive substantial diffusion over behaviorally relevant timescales. This issue should not be confused with another difficulty that arises in ring networks with few neurons, namely, that ring networks do not generically possess a true continuum of persistent states. Instead, they have a discrete set of stable fixed points, whose number is typically equal to the number of neurons. The latter problem was recently addressed in [33], where it was shown that recurrent weights can be tuned to rescue the continuity of the attractor. However, such weight tuning does not mitigate random diffusion (in fact, barriers between steady states diminish random diffusion, and their elimination enhances diffusivity).

Based on these considerations, it appears that small CANs in the brain must be structured in a way that allows for stabilization of the memory against ongoing neural noise, yet the computational principles that may underpin this stabilization are unknown. In the present work we identify a mechanism that can greatly reduce random diffusion in CANs, which is based on a combination of slow excitation and faster inhibition. We refer to this mechanism as *Negative Derivative Feedback* (NDF), following [45, 46] where it was shown, in a determinstic setting, that NDF can stabilize the dynamics of noise-free CANs to weight mistuning. However, it has been unknown whether NDF can reduce noise-driven diffusion in noisy neural networks.

NDF operates by using the derivative of a dynamical system’s readout in order to counterbalance undesired changes in its state (see Fig. 1B). The derivative signal can be generated within a neural network by utilizing a combination of slow excitatory and fast inhibitory synaptic currents (Fig. 1A, 1C). The sum of these currents cancels out when the readout is stationary, yet whenever the readout varies dynamically, it yields a non-vanishing signal, approximately proportional to the readout’s temporal derivative, that can serve as a feedback signal.

**Fig 1.**
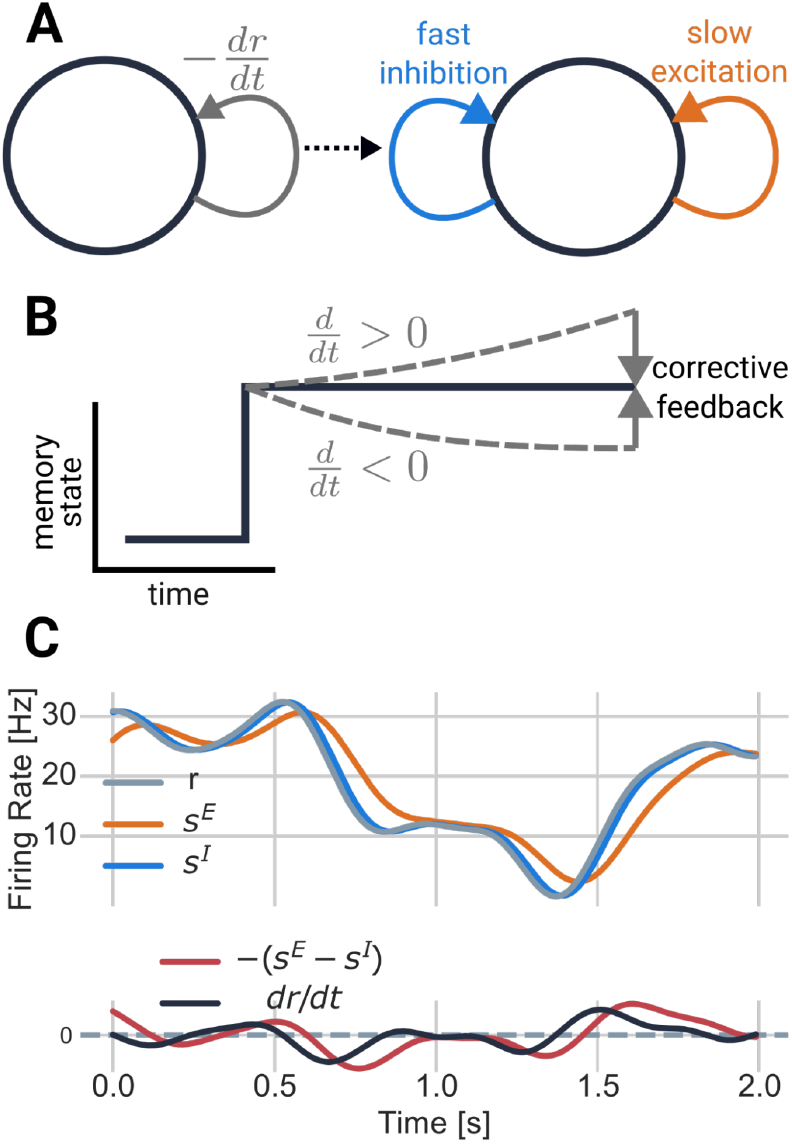
Principles of negative derivative feedback. **A**: Schematic illustration of a simple homogeneous network with negative derivative feedback. *Left* : neural network with idealized, explicit negative derivative feedback (see supplementary material, A.3). *Right* : negative derivative feedback is implemented approximately by a combination of slow excitatory and fast inhibitory synapses. **B**: The effect of negative derivative feedback on dynamics. When the network firing rate increases (decreases) from the initial memory state due to a perturbation, its derivative dr*/*dt becomes larger (smaller) than zero. NDF then acts in the opposite direction, correcting activity back towards the initial state. Panels A and B adapted from [45]. **C**: Approximating the derivative via synaptic dynamics. *Top*: Slow (excitatory) and fast (inhibitory) synaptic currents, denoted *s*^*E*^ and *s*^*I*^ represent a weighted average of the firing rate over its recent history, within a long (short) temporal window. *Bottom*: An approximation to the derivative is generated by subtracting the weighted average over a long interval from the weighted average over a short interval (see supplementary material, Eq. S14).

To analyze how NDF affects the dynamics of noisy neural networks, we first present a theory that analytically predicts diffusivity in a broad class of stochastic CANs with heterogeneous synaptic timescales, exhibiting excellent agreement with stochastic spiking simulations. We then show that when CANs are endowed with the NDF mechanism, it substantially mitigates the degradation of memory by noise-driven diffusion, and confirm this prediction in simulations of linear and nonlinear spiking CANs.

In key examples of CANs in the brain, steady state patterns span a low-dimensional *nonlinear* manifold. The principles of noise supression by NDF can be largely demonstrated and understood in simple linear neural networks, yet attempts to implement NDF in nonlinear networks such as the ring attractor introduce additional complexity. Due to the nonlinearity it is impossible to construct a linear readout of the activity that approximates the derivative of the stored memory, whose weights are invariant to the position along the attractor. As a result, we show that naive attempts to indiscriminately oppose changes in the neural activity do not only slow down motion along the attractor, but also slow down other fluctuations in the neural activity pattern, such as changes in the amplitude of the activity bump in a ring attractor.

We next show that even in this case it is possible to implement negative derivative feedback which is restricted only to the dynamics along the attractor, using the ring attractor as a prototypical example of a CAN with a nonlinear attractor manifold. This is achieved using a multi-layer architecture, in which the tasks of sensing motion and generating compensatory motion are carried out by distinct neural sub-populations. In this architecture diffusion is slowed down, and at the same time, neural activity remains tightly bound to a one dimensional ring manifold. A key insight is that some neurons previously thought to be only involved in velocity path integration may also take part in suppressing noise-driven diffusion. This can be of particular relevance in species in which the head direction network is structured as a ring attractor with a small number of neurons.

## Results

### Diffusion and continuous attractors

Individual units in neural networks are subjected to noise from various sources. As a consequence, some stochasticity is inevitable in any computation implemented by neurons. In neural networks that autogenously maintain short-term memory in their ongoing activity, stochasticity results in gradual corruption of the encoded information. This is especially true in networks that maintain persistent representations of continuous variables, since in such networks noise can easily shift the state of the network along the continuum of steady states [37, 40, 47].

Fig. 2 illustrates how memory degrades in a head-direction (HD) cell network. The network consists of cells tuned to distinct heading angles, with each cell firing most intensively when the head direction corresponds to its preferred angle. When cells are organized according to their preferred angle, a ring-like structure emerges, in which neighboring cells have similar orientation preferences (Fig. 2A). During spacial navigation, a bump of activity keeps track of an animal’s current heading (Figs. 2C-E). When the animal is still, the bump is persistently maintained in the location associated with its current heading, even in the absence of external stimuli (such as in total darkness [5, 18, 30, 48]). The memory state-space can thus be conceptualized as a ring (Figs. 2A-B).

**Fig 2.**
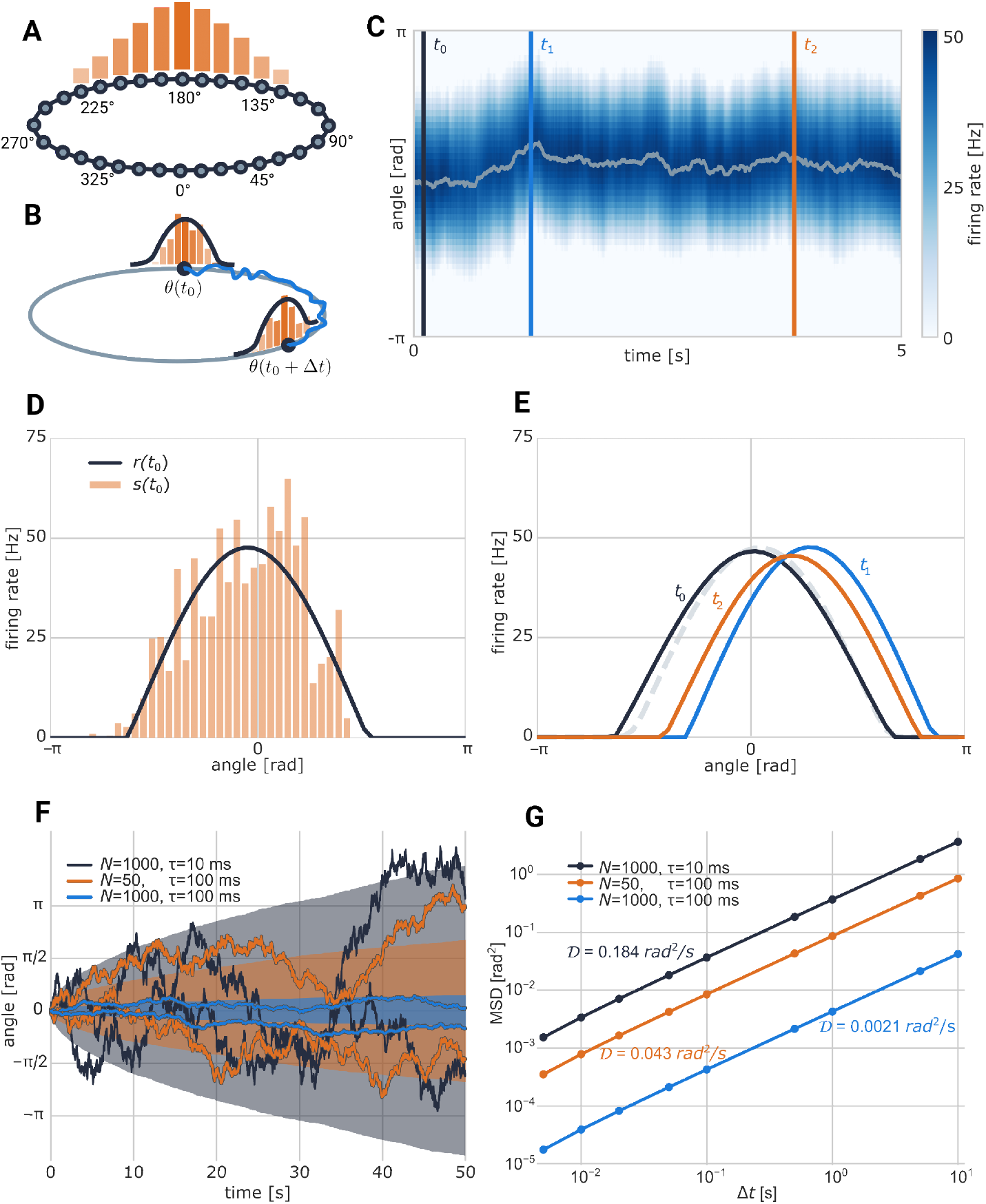
Continuous attractors and diffusion. **A**: Schematic of a ring attractor network. Bars indicate firing rates. **B**: Illustration of the noisy system’s traversal of the attractor over a time period Δ*t*, indicated by the blue line, with initial and terminal states marked by black circles. Noise causes distortions in the shape of the activity bump, which can be conceptualized as deviations from the attractor manifold within the *N* -dimensional space of neural activity. While most of these fluctuations decay rapidly and are therefore short-lived, noise also induces diffusive motion along the continuous attractor, which builds up over time. **C**: Activity of a simulated ring attractor network over 5 seconds. The x-axis indicates time while the y-axis indicates neuron index. Hue indicates firing rate. The gray trace marks the center of weight of the bump, which is also the readout angle *θ*. Vertical lines mark points in time at which snapshots of network activity are shown in panels D and E. **D**: Snapshot of firing rates and synaptic activations of the system at time *t*_1_. The synaptic activations are noisy due to the stochastic nature of the spiking activity. In turn, this noise induces fluctuations in the amplitude, width, and position of the firing rate profile. **E**: Snapshots of the firing rate profile at three points in time, color-matched to the vertical lines in panel C. Light gray dotted line shows average centered network activity. While bump width and amplitude exhibit limited fluctuations around their mean, fluctuations in its position accumulate over time. **F**: Three sets of trajectories of the angle readout from different different ring networks. *N* = 1000, *τ* = 10 ms (black), *N* = 1000, *τ* = 100 ms (blue), *N* = 50, *τ* = 100 ms (orange). Colored areas denote standard deviation. **G**: MSD-curves of the angle readout for the three networks in F. Referring back to panel A, MSD(Δt) is calculated over many trajectories of *θ* by taking their squared average ⟨ (*θ*(*t*_0_)− *θ*(*t*_0_ + Δ*t*))^2^⟩ . Diffusivity is normal as evidenced by linear MSD-curves, with logarithmic slope 1. The magnitude of the diffusivity is quantified by the diffusion coefficient 𝒟 (see Eq. 50).

In deterministic CAN models of the HD system [26–28, 33, 46, 49], short-term memory can be maintained indefinitely. However, when the stochastic nature of neural activity is taken into account, ring attractor networks exhibit diffusive random motion of the represented heading direction *θ*. Quantitively, diffusion can be characterized by its mean squared displacement (MSD) curve, which quantifies how far, on average, *θ* deviates from its starting point after some time Δ*t* has passed (Fig. 2B). If neural noise is uncorrelated in time, it induces *simple* diffusion of the stored memory, characterized by a MSD that grows linearly with Δ*t*. Diffusivity is then described by a single variable, the diffusion coefficient 𝒟, which measures the variance of drift in the represented variable per unit time (Fig. 2F).

Fig. 2F,G shows the MSD curves along with the diffusion coefficients of a simplified model of the head direction system, inspired by the one found in many insect species (model schematic in Fig. 2A). In this example, we take the time constants of excitation and inhibition to be identical. The time constant *τ* strongly affects diffusivity, with short synaptic time constants leading to rapid diffusion even in large networks (*N* = 1000). The insect head direction system stands out in this context because it contains only a small number of neurons [18, 30, 31, 33, 34, 44] – a feature which makes such systems especially prone to diffusion [40]. The orange plot in Fig. 2F shows the MSD for such a small network (*N* = 50).

Even when endowed with a relatively long synaptic time constant (*τ* =100 ms), the network with 50 neurons exhibits strong diffusion corresponding to average root mean square displacement of more than 15 degrees over the course of a second. Furthermore, while in this work we focus on the noise generated by the neurons within the attractor network itself, neurons in biological neural circuits are subjected to additional noise via inputs from other brain circuits, which is expected to further contribute to the diffusivity.

Considering the exquisite ability of some insect species to maintain short term memory of heading and position using remarkably small neural circuits [18], these observations raise the question, whether there are mechanisms that can mitigate noise driven diffusion in ring attractors and in CANs with other network architectures. Next, we examine the consequences of NDF in noisy neural networks, and show that this mechanism can dramatically reduce diffusion.

### Line attractor network

We start by examining the effect of NDF on diffusivity in the line attractor, which is the simplest attractor topology that can be supported by a CAN. Despite their relative simplicity, line attractors are hypothesized to support various functions in the brain, such as the control of eye position by the oculomotor integrator [10–13, 43] and decision making [50, 51]. We focus here on a simple neural architecture with linear neural response functions, allowing us to gain precise mathematical insights, which we later generalize to a much broader class of CANs.

The simplest neural network architecture that supports a line attractor consists of a single homogeneous population of *N* neurons, which contact each other with unstructured excitatory and inhibitory connections. Fig. 3 illustrates the architecture of such a network, in which the excitatory and inhibitory synaptic connection strengths are parametrized by *J*_*E*_ and *J*_*I*_. In order to allow for NDF, excitatory and inhibitory synapses are modeled as having distinct timescales, *τ*_*E*_ and *τ*_*I*_. Additionally, the population can receive an external excitatory input.

**Fig 3.**
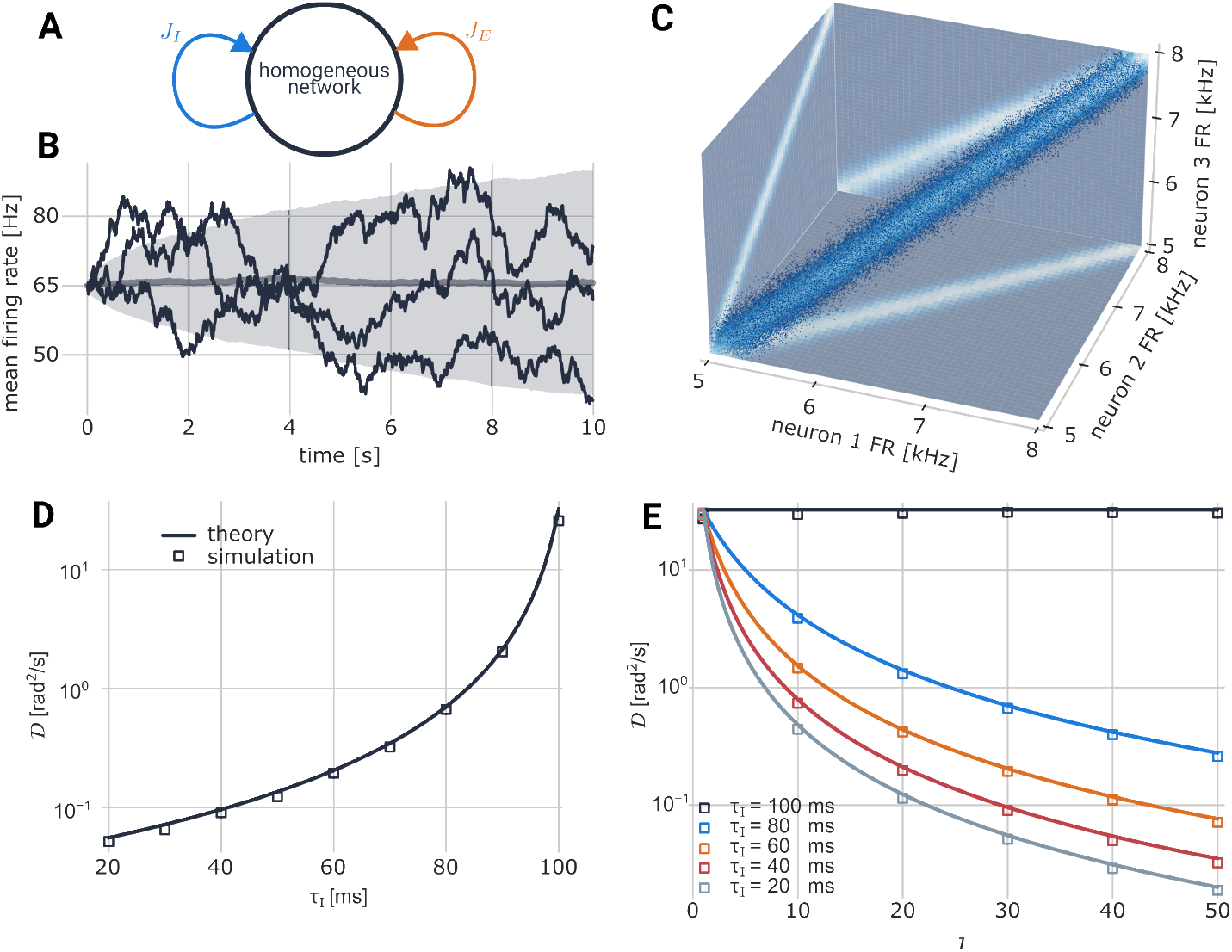
Diffusion in the line attractor network. **A**: Network schematic. Excitatory (inhibitory) connections strengths are denoted by *J*_*E*_(*J*_*I*_). **B**: Example trajectories of network activity (mean firing rate as a function of time, black traces). Thick gray trace: mean trajectory across many realizations. Shaded area (in light gray): standard deviation. **C**: Visualization of the attractor. Blue dots depict firing rates (spike trains convolved with an exponential kernel with a 100 ms time constant), sampled at dense time points across 500 simulations of duration 10 s each with a uniform distribution of initial values between 4 and 9 kHz. Color of dots: joint probability of all three firing rates; coloring of the surfaces: pairwise joint probability for each pair of neurons (lighter colors represent higher probability). Here, *τ*_*E*_ = 100 ms and *τ*_*I*_ = 50 ms. **D**: Diffusivity as a function of the inhibitory timescale *τ*_*I*_ while the excitatory timescale *τ*_*E*_=100 ms is kept constant. *J* = 30 (solid trace: theory, square symbols: simulations). **E**: Diffusivity as a function of the feedback gain parameter *J*, shown for different values of *τ*_*I*_ (solid traces: theory, square symbols: simulations).

Because of the homogeneous connectivity, the *N* dimensional dynamics can be reduced to a description that involves two variables: the average, over the whole neural population, of excitatory synaptic activations, *s*^*E*^, and an average across the population of inhibitory synaptic activations, *s*^*I*^. In order for NDF to arise, we require that *τ*_*I*_ ≤ *τ*_*E*_.

When *J*_*I*_ = *J*_*E*_ − 1, the network has a continuum of steady states, which form a line attractor (see supplementary material, A). Under this condition the network can sustain persistent activity at any firing rate in the absence of external inputs through its reverberating activity. The connection strengths that yield a line attractor can be parameterized using a single connectivity parameter *J*, such that:

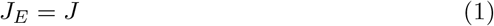

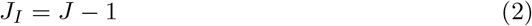

The parameter *J* governs the strength of both inhibition and excitation. Importantly, it does not affect the net synaptic input at steady state: excitatory and inhibitory inputs both increase with *J*, precisely canceling each other when neural activity is stationary. However, when neural activity varies over time, the difference in excitatory and inhibitory timescales produces a transient signal which approximates the derivative of the firing rate, Fig. 1. Hence, *J* represents the gain of negative derivative feedback.

We model the system as consisting of Poisson-spiking neurons (see Methods, *Network Model*), thereby introducing stochasticity into the dynamics. Our goal is to examine whether noise resilience is enhanced by NDF, and how this is affected by feedback gain and the synaptic timescale difference. For simplicity, we keep *τ*_*E*_ fixed at 100 ms and vary *τ*_*I*_ as the sole timescale parameter.

Due to the spiking noise, the firing rate exhibits diffusive drift along the line attractor. The synaptic activities, *s*^*E*^and *s*^*I*^, vary along a line as shown in Fig. 3C. In networks with a single synaptic time scale *τ*, diffusivity scales as *τ* ^−2^ [40] and therefore increases with decrease of *τ*. However, negative derivative feedback has an opposite effect: as the timescale of inhibition shrinks, while the timescale of excitation remains constant, the diffusion coefficient *decreases*, Fig. 3D, while steady-state dynamics are not affected. This result is striking because, naively, we associate a shortening of synaptic timescales with faster dynamics, less persistence and therefore higher diffusivity. Similarly, increasing the feedback gain decreases diffusion as well, Fig. 3E.

Further theoretical analysis (see supplementary material, D) provides deeper insights on how the diffusion coefficient scales with the synaptic timescales *τ*_*E*_, *τ*_*I*_ and the connectivity parameter *J*, with the following expression being derived from Eq. S89:

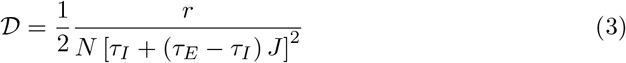

where *r* is the average firing rate of any given neuron in the network. The accuracy of this expression is validated in Fig. 3D-E. Diffusivity decreases linearly with the number of neurons *N*, since redundancy helps average out noise.

When synaptic timescales are equal (*τ*_*E*_ = *τ*_*I*_ = *τ*), diffusivity scales as *τ* ^−2^, as shown in previous works. Moreover, in this case, the feedback gain *J* does not affect diffusivity at all. We see, however, from Eq. 3 that decreasing *τ*_*I*_, while keeping *τ*_*E*_ fixed, always reduces diffusivity. This prediction is confirmed by simulations (Fig. 3D-E). Furthermore, the feedback gain *J*, which has no influence on the steady states of the system, does affect its stochastic dynamics. As long as *τ*_*E*_ *> τ*_*I*_, diffusivity is a monotonically decreasing function of *J* (scaling asymptotically as *J*^−2^ for large *J*, Eq. 3). This prediction, too, is quantitatively validated by the simulations, Fig. 3E.

### A theory of diffusivity in nonlinear CANs

The previous result for line attractor networks can be generalized to a much larger class of models, namely all CANs made up of Poisson spiking neurons with an arbitrary number of synaptic timescales.

The most general expression is derived and discussed in Methods, and here we focus on networks with two synaptic timescales, excitatory (*τ*_*E*_) and inhibitory (*τ*_*I*_), since this is sufficient to implement NDF. In this case, the analytical expression for the diffusivity takes the form

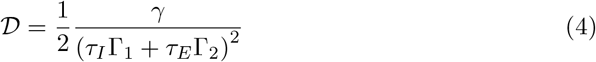

where *γ*, Γ_1_, and Γ_2_ depend on the architecture of the network (see Eq. 56), but not on the timescales. The key quantities that determine these coefficients are the sensitivities 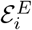 and 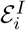 (Eq. 57), which quantify the susceptibility of the excitatory and inhibitory synaptic inputs impinging on neuron *i* to changes in position along the attractor.

In methods we show that under fairly unrestrictive requirements, Γ_1_ ≤0 and Γ_2_ ≥0, which implies that diffusivity is decreased by reducing the timescale of inhibition. In particular this is true if for all active neurons, two conditions are met: first, that 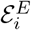 and 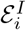 have opposite signs. This means that as the attractor is traversed, an increase (decrease) in excitation is accompanied by an increase (decrease) in inhibition: in other words, changes in excitatory inputs are balanced by changes in inhibitory inputs. The second condition is that 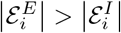,which means that excitation is more sensitive to changes in attractor position than inhibition.

The first condition in particular highlights a hallmark of NDF, namely, that fast inhibition is balanced out by slow excitation. Thus, the components of the synaptic connectivity responsible for NDF are expected to produce covarying excitatory and inhibitory synaptic inputs. The remaining components of the connectivity are responsible for the excitatory and inhibitory synaptic inputs that sustain the network’s steady states. In summary, under a broad range of requirements, diffusivity is decreased by reducing the time scale of inhibition. Furthermore, these requirements are compatible with the key characteristics of NDF: balance between slow inhibition and fast excitation.

### The ring attractor network

Next, we demonstrate how NDF can effectively reduce diffusion in a concrete example of a nonlinear CAN architecture, namely the ring attractor network. As discussed in previous sections, the ring attractor network is often used to model the head direction system of mammals and insects [26–29, 33], with extensive support from experimental evidence [15, 18, 30, 31, 34, 44].

We focus on a recurrently connected network model consisting of a small set of *N* Poisson spiking neurons arranged on a ring, Fig. 4A. Neurons form slow excitatory and fast inhibitory connections with each other, with both excitation and inhibition being strongest between neurons with similar orientation, Fig. 4B. More precisely, connectivity is given by

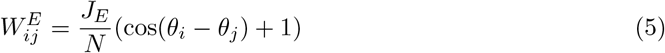

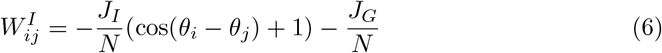

where *θ*_*i*_ is the preferred firing angle of neuron *i*, and the excitatory (inhibitory) synaptic activation of neuron 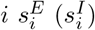 is governed by the timescale *τ*_*E*_ (*τ*_*I*_), with which they track the neuron’s firing rate *r*_*i*_. Biological data supports the plausibility of a similar excitatory connection scheme in multiple species of insects [30, 31, 34, 44, 52], and a similar inhibitory connection scheme in at least one [53, 54]. The Cosine dependence of the weights on *θ*_*i*_− *θ*_*j*_ was chosen for simplicity, but all considerations in this section readily extend to similar symmetric connectivity profiles.

**Fig 4.**
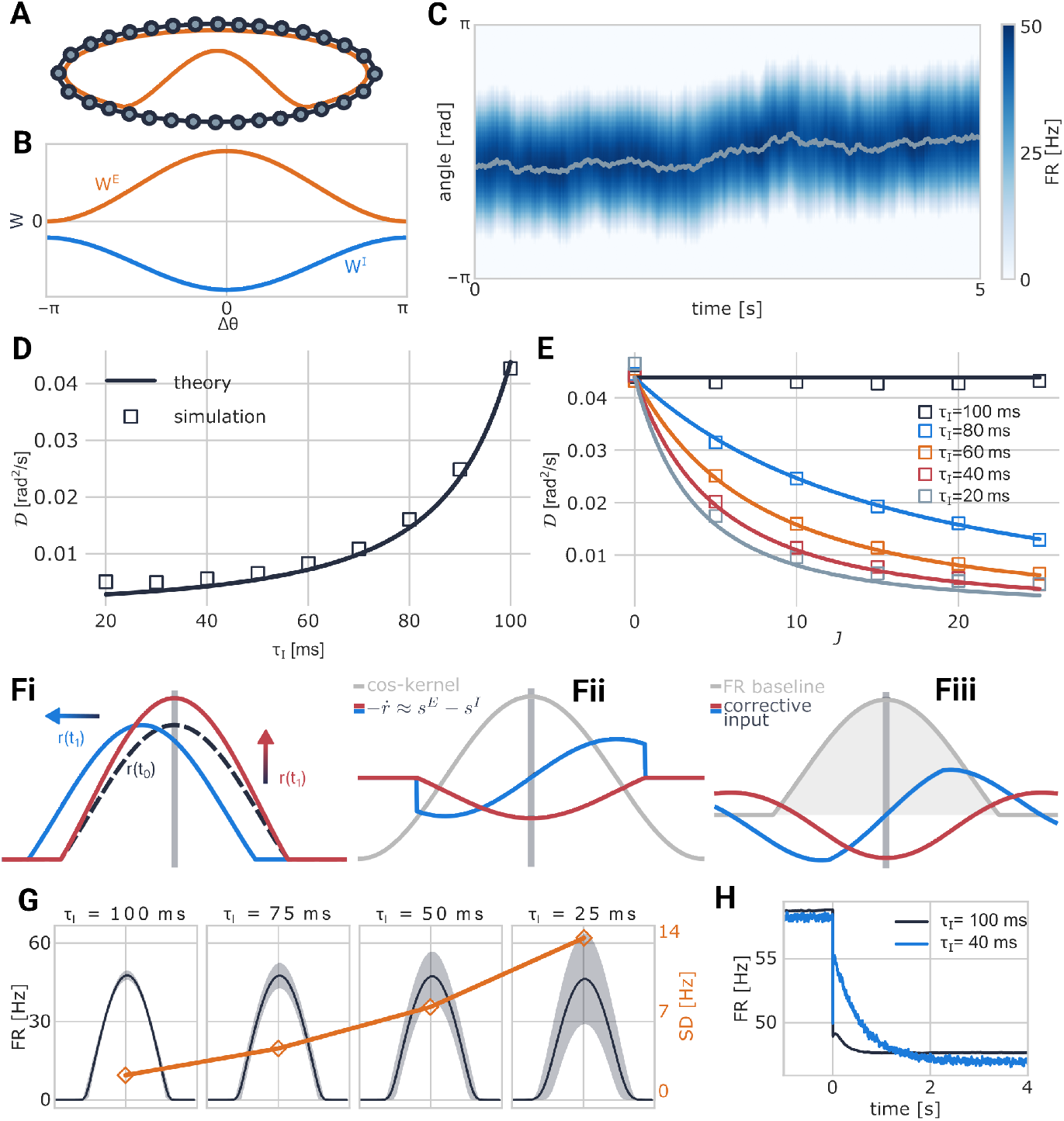
The ring model. **A**: Network schematic. Neurons are arranged on a ring, and connectivity depends on the distance between neurons as shown in B (orange trace: example of neural firing rates at steady state). **B**: Network connectivity as a function of distance on the ring. Both excitation and inhibition are stronger between neurons with similar orientation (see Eqs. 5-6). **C**: Example network activity. Gray trace: angle readout. **D**: Diffusivity as a function of the inhibitory timescale *τ*_*I*_ while the excitatory timescale *τ*_*E*_=100 ms is kept constant. *J* = 20 (solid trace: theory, square symbols: simulations). **E**: Diffusivity as a function of the feedback gain parameter *J*, shown for different values of *τ*_*I*_ (solid traces: theory, square symbols: simulations). **F**: Illustration of the corrective mechanism in the ring network. *i* : Two examples of possible perturbations of a bump state (dashed trace): shift along the attractor (blue trace) and amplitude perturbation (red trace). *ii* : temporal derivatives of the firing rates associated with the perturbations shown in *i* (blue and orange traces). These are approximated by *s*^*E*^ − *s*^*I*^ and are subsequently convolved with a cosine shaped kernel (gray trace, Eq. 21) to generate the feedback signal. *iii* : The resulting feedback signal comprises an input to the neurons that opposes the perturbation by inducing an opposing shift in position (blue trace) or change in amplitude (red trace). **G**: NDF causes increased fluctuations in bump amplitude. For *τ*_*I*_ ∈ [100, 75, 50, 25] ms (from left to right). Black trace: average centered bump. Blue area: fluctuations in centered bump amplitude ranging between the 10th and 90th percentiles. Orange: standard deviation of firing rate at the bump center. **H**: Relaxation of bump amplitude after a perturbation. The network is stimulated up to time *t* = 0 with an additional, spatially modulated input that increases the amplitude of the bump. At *t* = 0 this input is removed and the amplitude relaxes back to baseline. Black trace: *τ*_*I*_ = 100 ms, black trace: *τ*_*I*_ = 40 ms. Averages over 5000 simulations.

Under certain conditions (see Methods, Eqs. 67-70), the stationary states of the dynamics are localized activity bumps, which can be centered around any position on the ring (see Eqs. S18 S28, S31, S33). Bump formation is driven by excitation between nearby neurons and inhibition between faraway neurons, which requires *J*_*E*_ *> J*_*I*_ (Eq. 69).

The synaptic input *g*_*i*_ to each neuron *i* can be split into two components: *g*_*i*_ = *G*_*i*_ + *δG*_*i*_ (see Methods, Eqs. 21-22). The first component, *G*_*i*_, is a spatially structured input which maintains the bump. The other component, *δG*_*i*_, is responsible for correcting network activity in response to perturbations. This term is a linear combination of differences in synaptic activations 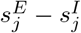,which approximate the negative derivative of the firing rates *r*_*j*_. Hence *δG*_*i*_ vanishes if the neural activity is stationary, and it acts as a corrective signal whenever the neural activity varies in time. More precisely, *δG*_*i*_ can be interpreted as implementing NDF on the bump location and amplitude, via low-order Fourier components (see Methods and [46]).

It is straightforward to see that the two conditions for Γ_1_ *<* 0, discussed in the previous section, are met for the network architecture discussed above (Methods). Consequently, it is guaranteed that introducing NDF into the dynamics by making *τ*_*I*_ *< τ*_*E*_ will reduce diffusivity, see also Eq. S105.

Numerical simulations confirm these theoretical considerations, and further demonstrate that the reduction in diffusivity can be very substantial. As in the linear network, the smaller *τ*_*I*_ is relative to *τ*_*E*_, the greater is the reduction in diffusivity, Fig. 4D. To highlight the similarity with the linear network, we parametrize the synaptic weights as 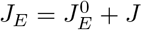 and *J*_*I*_ = *J*, where *J* controls the gain of negative derivative feedback, i.e., the magnitude of *δG*_*i*_. The parameter *J* has no effect on the steady states of the network (see Methods), yet increasing it can dramatically reduce diffusivity, provided that *τ*_*E*_ *> τ*_*I*_, Fig. 4E, see also S108. Thus, *J* plays a similar role as the gain parameter in the linear network.

We note that the analytical theory tends to slightly underestimate diffusivity when the timescale difference is large, especially under strong feedback gain (compared to simulations, see Fig. 4D-E). The mismatch is due to large fluctuations in bump amplitude that are introduced by NDF under these conditions. The theory assumes small deviations between the synaptic activations and their steady state values, and it therefore becomes less accurate when the deviations are large.

### Naive application of NDF disrupts attractor tightness

Alongside its success in suppressing noise-driven diffusion, the feedback mechanism also introduces some dynamical artifacts, which are potentially undesirable. In the model introduced above, NDF does not specifically target movements of the bump (Fig. 4F). Consequently, perturbations of the bump amplitude relax more slowly than in the absence of NDF (Fig. 4H). In addition, the timescale difference between strong excitatory and inhibitory inputs associated with NDF induces firing rate fluctuations, which manifest as large amplitude variations (Fig. 4G). Conceptually, the fluctuations in the amplitude of the bump imply that network activity is not tightly bound to the ring attractor. However, there are examples or neural representations that are strictly bound to a low-dimensional manifold, despite dynamic fluctuations in the represented variable [15, 55]. These dynamical artifacts raise the question whether negative feedback can suppress noise-driven diffusion *without* slowing down perturbations affecting the shape of the bump, thereby retaining activity which is tightly bound to a low-dimensional attractor.

### Precise negative derivative feedback in a multi-layered architecture

It is instructive to think of the NDF mechanism as composed of two separate computations: a measurement of bump velocity, extracted from the neural activity, and induction of compensatory motion in response to this measurement. To maintain attractor tightness, the induction of compensatory motion must not affect the bump in any way other than a shift in its position. Likewise, the extracted measurement should be sensitive to the velocity of the bump, while remaining insensitive to other deformations. In other words, it should yield a signal that allows specifically for NDF of bump motion.

We start with elucidating how the bump velocity measurement can be extracted from the neural activity. As above, 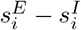 approximates the derivative of the firing rate, 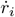. If the bump position is known, it is straightforward to construct a linear combination of these quantities that corresponds to an estimate of the velocity. However, the appropriate linear combination depends on the position of the bump (see supplementary material, C.1). Consequently, measurement of bump velocity must involve a nonlinear computation.

The required computation can be implemented using a multi-layer structure (Fig. 5A). The network includes a layer of recurrently connected neurons which are responsible for sustaining a localized activity bump (population *C*). Another set of neurons (populations *V*_*L*_ and *V*_*R*_) are dedicated to the extraction of velocity, and are driven by activity in the central bump. We refer to these neurons as velocity-sensing neurons. Here we briefly discuss the connectivity of these neurons, and additional details are given in Methods, *Multi-Ring Model*.

**Fig 5.**
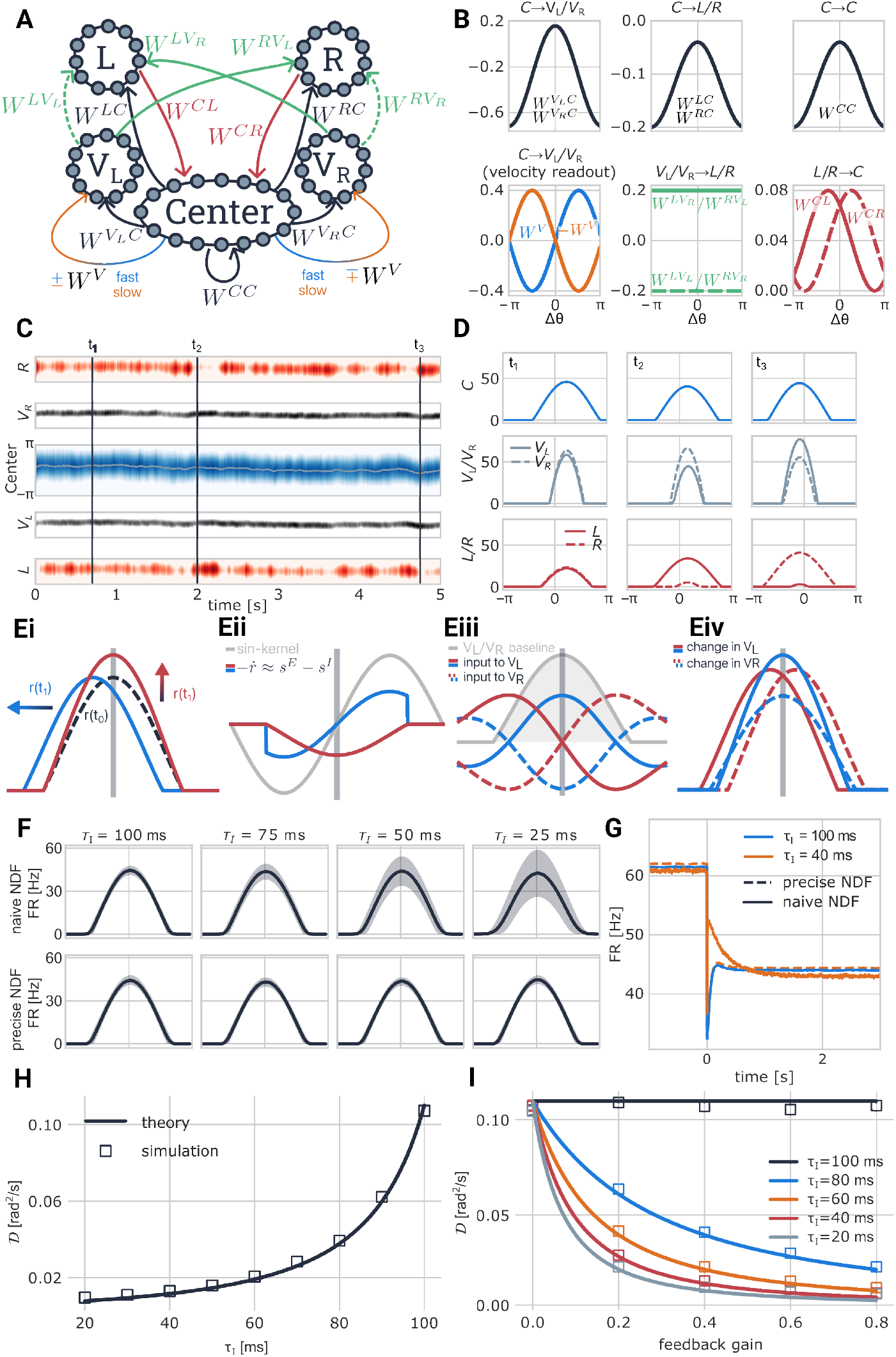
The velocity-feedback model. **A**: Network schematic. Pathways along which the velocity feedback is propagated are colored, while all other connections are black. In addition to the the central ring *C, V*_*L*_, *V*_*R*_ are populations of velocity-sensing neurons, L,R are populations of rotation cells. **B**: Visualization of all connections in A (synaptic weights as a function of preferred angle difference). The central ring connects to the *V*_*L*_ and *V*_*R*_ populations through three types of synapses: symmetric-weight synapses (black), which induce an inherited bump in the *V*_*L*_ and *V*_*R*_ populations, and asymmetric-weight synapses – fast (orange) and slow (blue) – which enable local velocity sensing. *L*/*R* connect to *C* with shifted cosine connections, enabling velocity integration (red). *V*_*L*_/*V*_*R*_ connect to *L*/*R* with all-to-all connectivity, compensating for random motion of the central bump (green). **C**: Example of network activity in all five rings over five seconds of simulation time. Times for which snapshots of activity are shown in D are marked with vertical lines. **D**: Snapshots of network activity at times *t*_1_, *t*_2_, *t*_3_. Blue represents activity of the central ring, gray activity in *V*_*L*_(solid) and *V*_*R*_ (dashed), red activity in *L* (solid) and *R* (dashed). In the left column (*t*_1_), the central bump has been fairly stationary in the past 100 ms. In the middle column (*t*_2_), it has been moving counter-clockwise. As a result, the *V*_*L*_ population is more active than the *V*_*R*_ population. This, in turn, is causing higher activity in the *R* population of rotation cells than in the *L* population, which will induce compensatory motion of the central bump to the right. The right column shows an example in which the central bump has been recently moving clockwise. **E**: Illustration of the corrective mechanism in the ring network. *i* : Two examples for possible perturbations of a bump state (dashed trace): shift along the attractor (blue trace) and amplitude perturbation (red trace). *ii* : temporal derivatives of the firing rates associated with the perturbations shown in *i* (blue and red traces). These are approximated by *s*^*E*^ − *s*^*I*^ and are subsequently convolved with a sine shaped kernel (gray trace, Eq. S S47) to generate the input to the velocity sensing neurons. *iii* : Inputs to the velocity sensing neurons *V*_*L*_*/V*_*R*_, in response to amplitude change in the central ring (red) and in response to counter-clockwise shift (blue). Solid traces: inputs to *V*_*L*_. Dashed traces: inputs to *V*_*R*_. Gray trace: baseline *V*_*L*_/*V*_*R*_ activity for stationary bump. *iv* : Resulting firing rates of rotation cells neurons *V*_*L*_*/V*_*R*_ in response to amplitude change in the central ring (red) and in response to counter-clockwise shift (blue). Solid (dashes) traces: firing rates of *V*_*L*_(*V*_*R*_). *L* and *R* receive input from *V*_*L*_ and *V*_*R*_ that is proportional to the difference of their total population activities (see Eqs. 41-44). This results in zero input in case of an amplitude change, since both populations are equally active (solid vs. dashed red trace), whereas in the case of a counter-clockwise shift, this results in induction of clockwise motion, since the total input to *V*_*R*_ is larger than the total input to *V*_*L*_ (blue vs dashed blue traces). **F**: Attractor tightness: variation of firing rates of the central ring. Black: mean activity. Gray shaded area represents the range of firing rates between the 0.1 and 0.9 quantiles of activity in *C*. Top row: naive NDF mechanism on central ring (see Methods). Bottom row: precise NDF mechanism as described in A. Left to right: decreasing *τ*_*I*_. **G**: Relaxation of bump amplitude following abrupt change in input to the central ring, at time 0. Relaxation is markedly slowed down under naive NDF (blue trace: *τ*_*I*_ = *τ*_*E*_ = 100 ms, no NDF; orange trace *τ*_*I*_ = 40 ms *< τ*_*E*_). Under the precise NDF scheme, relaxation of bump amplitude is nearly unaffected by NDF. **H**: Diffusion coefficient as a function of *τ*_*I*_, for the precise NDF architecture. Feedback gain is set to *v*_*g*_ = 0.4 (see Methods *Parameters*). **I**: Diffusion coefficient as a function of the velocity feedback gain.

Velocity sensing neurons are driven by activity in the central bump. Their connectivity can be split into two components: the first (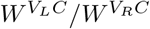 in Fig. 5A-B), is structured in a similar way as the recurrent connectivity within the central population, causing *V*_*L*_ and *V*_*R*_ neurons to inherit a bump of activity from the central population, that moves in phase with the central bump. Second, each velocity-sensing neuron in *V*_*R*_ (*V*_*L*_) is driven by a linear combination of the derivative estimates *s*^*E*^ − *s*^*I*^. The linear combination projecting to neuron *i* in the *V*_*L*_ (*V*_*R*_) population is designed to accurately extract the velocity of the central bump moving in the clockwise (counter-clockwise) direction, provided that the bump is centered around *θ*_*i*_. This is achieved by a projection on a sinusoidal pattern (*W* ^*V*^), centered around *θ*_*i*_, bottom left panel in Fig. 5B (a variant of the connectivity pattern, in which weights are purely excitatory or purely inhibitory, is presented in the supplementary material C.3, and yields very similar results).

While a neuron precisely at the center of the bump correctly extracts the velocity, the neuron on the opposite side of the ring measures the same velocity but in the opposite direction, and all other neurons measure something in between these two extremes. However, due to the additional bump-inducing input from the central ring, this velocity signal modulates only the activity of neurons whose preferred orientation is close to the current bump location. As a consequence, neurons in the *V*_*R*_ population exhibit increased (decreased) firing rates when the bump moves clockwise (counter-clockwise), and the *V*_*L*_ population exhibits opposing behavior. Overall, the summed activity of all *V*_*R*_ neurons, minus the summed activity of all *V*_*L*_ neurons, generates a precise measurement of velocity (see supplementary material, C.2). Fig. 5E illustrates how velocity is read out by this machinery and how it is affected by shifts in orientation but not amplitude.

We have now discussed how to extract a measurement of bump velocity, but not yet addressed how it can be fed back into the network in a way that corrects perturbations in bump position. To that end, we introduce another set of neural populations, which we collectively refer to as rotation cells (following [26]). They comprise the populations *L* and *R*, which inherit the bump from *C* (via connections *W* ^*LC*^/*W* ^*RC*^) and project back to the center with shifted connectivity (see Fig. 5A-B). Populations with similar connectivity are often included in models of head-direction cells in order to account for velocity integration [10, 26–28, 33] and robust evidence for their existence has been found in the fly [18, 30, 31, 34, 44, 56, 57]. Stimulating *R* (*L*) shifts the central bump (counter-) clockwise in the center, and all other populations follow suit.

It is now easy to see how the activity of *V*_*L*_ and *V*_*R*_ can be harnessed to provide an effective feedback signal: we introduce uniform, positive connectivity from *V*_*L*_ to *L* and negative uniform connectivity from *V*_*R*_ to *L*, and vice versa for the *R*-population. When noise drives motion of the central bump, the *L* and *R* populations are differentially driven by *V*_*L*_ and *V*_*R*_, causing compensatory motion of the bump in the opposite direction.

Figs. 5I-J demonstrate that in the above architecture, diffusion is mitigated by the NDF mechanism, as observed above in other architectures. At the same time, the NDF mechanism does not induce amplitude fluctuations, in contrast to what is seen in the simpler architecture (Fig. 5G). In addition, the relaxation of amplitude perturbations is not slowed down by the NDF either (Fig. 5H). Since the structure of the bump remains highly stereotyped, neural activity in the central bump remains tightly bound to a one-dimensional manifold (Fig.5G).

Finally, in order to compare the two implementations of NDF within the same multi-layer architecture, we consider a variant of the connectivity, where we replace the precise NDF signal (by setting *W* ^*V*^ to zero) with a naive NDF mechanism akin to the one from the previous section: the center population projects slow excitation and fast inhibition onto itself (see Eqs. 33, 48-49), and the feedback gain is regulated by the strengths of these connections (see previous section). With this setup, it is easy to see that without NDF, both networks will exhibit the same diffusivity. We then set the feedback gains such that the dependence of diffusivity on synaptic timescale matches in the two variants of the network (see Fig. S5). With this matching, naive NDF introduces much larger amplitude fluctuations than the precise NDF mechanism (Fig. 5G). In addition, naive NDF slows down the relaxation of amplitude fluctuations, which is not observed when feedback is limited to bump velocity (Fig. 5H). In conclusion, precise NDF yields a similar reduction in diffusivity as simple NDF, while tightly confining the activity in the central bump to a one dimensional manifold.

## Discussion

We showed that negative derivative feedback can effectively mitigate noise-driven diffusion in continuous attractor networks. Using a theory that accurately predicts diffusivity in Poisson spiking CANs with multiple synaptic timescales (see also [41]), we identified a set of sufficient conditions for reduction in the diffusivity due to time-scale heterogeneity. These conditions encompass the network architectures proposed in [45] for stabilization of the attractor to weight mistuning. Hence, we identified organizational features of the synaptic connectivity that simultaneously stabilize CANs to drift arising from dynamic and structural sources of noise.

In networks employing NDF, an increase in excitatory input to each neuron is closely mirrored by a corresponding rise in inhibitory input. At steady state, particularly under conditions of high NDF gain, inhibitory input effectively balances the majority of excitatory input. This dynamic aligns with the well-established balance of excitation and inhibition, a hallmark of cortical activity [45, 58–62]. Additionally, the stability of continuous attractor networks (CANs) can be further reinforced by incorporating short-term synaptic plasticity mechanisms [63–65].

Diffusive dynamics of the memory stored in CANs is predicted based on theoretical considerations, yet it has also been observed in recordings of neural activity from brain areas hypothesized to implement CAN dynamics. Two important examples are the oculomotor integrator network, where diffusion has been linked to fixational eye drifts [43], and the mammalian head direction system [15]. Diffusive motion along the continuous attractor may be beneficial in the oculomotor system [66], but it is likely detrimental for the maintenance of short term memory in the head direction system.

Extensive knowledge has been gained in recent years on the dynamics and architecture of the head direction system in several insect species. Even though it is premature to seek direct parallels between the architecture of the insect ring attractors and the models studied here, it is intriguing to note that several of their features may be conducive to NDF. In the ellipsoid body of the desert locust, a species capable of traveling large distances through fairly featureless terrain, similar tuning has been observed in excitatory and inhibitory connections between the compass neurons [52–54, 67, 68]. If inhibitory time constants are shorter than the excitatory ones, it is likely that the conditions for naive NDF are met.

The precise NDF architecture (Fig. 5A) involves several sub-networks with distinct anatomical and functional properties. All sub-networks express a bump of activity that represents heading, yet in some sub-networks neurons are also selective to motion. Such bump inheritance is a ubiquitous feature across various populations in the fly head direction system [31, 34]. Various cell types are co-tuned to angular velocity and head direction [30, 31, 34, 56], a characteristic feature of the *V*_*L*_ and *V*_*R*_, as well as the *L* and *R* populations in our model. Furthermore, some of the co-tuned cells in the central complex have been demonstrated to facilitate velocity integration by inducing activity shifts via connectivity similar to *W* ^*CL*^ and *W* ^*CR*^. The *L* and *R* neurons in our model were modeled in accordance with these properties. We thus hypothesize that neurons involved in velocity integration, such as the PEN-1-cells in the fly, could also be implementing NDF by inducing compensatory motion in response to internal drifts. A prediction arising from this hypothesis is that even when external inputs are absent, candidate velocity sensing and rotation neurons will exhibit selectivity to motion of the central bump, generated either internally due to noise, or via optogenetic stimulation of the E-PG neurons. Finally, it is important to note that the multi-ring network (Fig. 5A) serves as a proof of principle that precise NDF can be attained, and variations on its architecture may achieve similar functionality.

## Methods

### Network model

#### Poisson model

We consider, in all parts of this work, networks consisting of *N* Poisson spiking neurons. The instantaneous firing rate of neuron *i* is given by

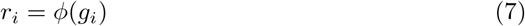

where *ϕ* is a nonlinear transfer function and *g*_*i*_ is the total synaptic input to neuron *i*. In all the specific examples analyzed in this work, *ϕ* is chosen as a ReLu function:

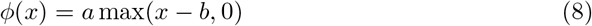

where the parameters are specified below (*Parameters*). Analytical calculations derived in this work are valid for any smooth, monotonic transfer function.

Each two neural units in the network are connected by a number of synapses with up to *M* distinct timescales *τ*_1_, …, *τ*_*M*_. The synaptic dynamics are given by

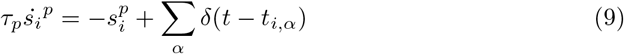

where *i* is the neural index, *p* is the index of synaptic timescale, and 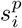 is the synaptic output with timescale *p* generated by the spikes of neuron *i* (up to the multiplication by the synaptic weight). The spike train of neuron *i* is represented by 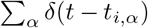,where *t*_*i,α*_ is the *α*-th spike time of neuron *i*. Finally, we define the input to neuron *i*, arising from synapses with timescale *p*, as

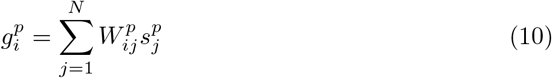

and the total synaptic input *g*_*i*_ to neuron *i* is given by

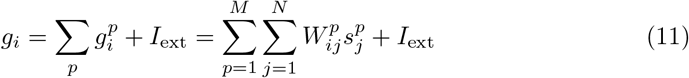

where *I*_ext_ represents an external synaptic current.

#### Rate model

We also consider a rate approximation of the Poisson spiking model, defined as follows. Equations 7-10 remain unaltered, except for equation 9, which describes the synaptic activation of neuron *i* with timescale *p*. This equation is replaced by

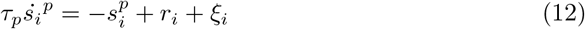

where *ξ*_*i*_ is a Gaussian noise term which approximates the fluctuations arising from the irregular Poisson spiking. Its statistics are given by

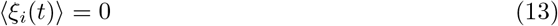

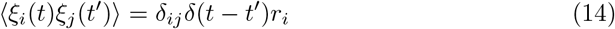

reflecting the fact that all neurons in the network receive independent noise with a variance proportional to their firing rate, due to the Poisson statistics. The Gaussian approximation becomes precise in the limit in which firing rates are sufficiently large, such that the number of inputs spikes driving each synapse, within its synaptic timescale is large (see [40]).

#### The attractor manifold

We assume that the connectivity, as defined by the matrices *W* ^*p*^, *p*∈ {1, …, *M*}, is constrained such that the network has a continuous attractor. This means that the deterministic dynamics, obtained by eliminating the noise term *ξ*_*i*_ in Eq. 12, are characterized by a continuum of steady states (at least in the limit *N* → ∞), which form a low-dimensional manifold within the *N* dimensional neural activity space (the attractor manifold). These steady states are parametrized by a variable *θ* and are denoted by 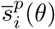.

#### Linear model

The dynamics of the linear network with slow excitatory and fast inhibitory synapses are specified by

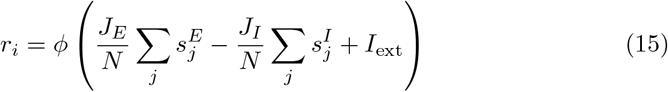

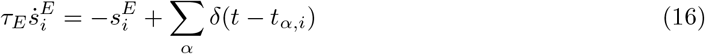

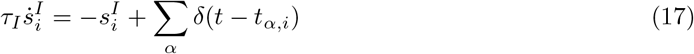

where *J*_*I*_ = *J*_*E*_− 1. For simplicity, autapses were not eliminated from the network, however, in a large enough network this has no effect on its dynamics. For the same reason, we do not enforce Dale’s law. It would be straightforward to split the network into excitatory and inhibitory populations without altering the key conclusions.

We are mainly interested in the dynamics of this network when *I*_ext_ = 0. The deterministic dynamics are then characterized by a continuum of steady states, representing memories that the network can retain in its persistent activity. A transient external current can be used to set the network firing rate to a desired value.

Because the network is homogeneous, it is equivalent to a network with a single neuron (*N* = 1), and its state can be precisely described by two dynamics variables *s*^*E*^ and *s*^*I*^ (see Supplementary Material A).

#### Ring model

The dynamics of the ring model are given by

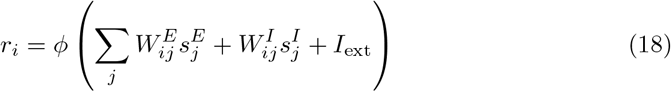

where 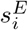 and 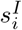 follow equations 16-17, like for the linear model. The connectivity is defined as follows:

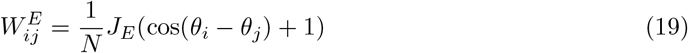

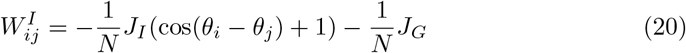

#### E-I balance in the ring model

In the ring model, the total input to neuron *i* can be written, using equations 11, 18-20, as

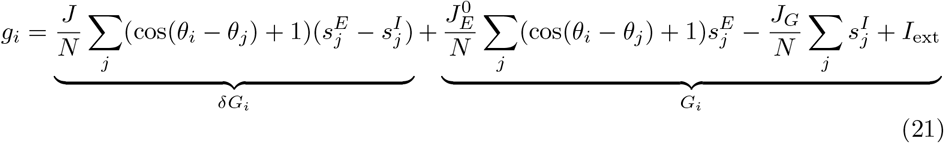

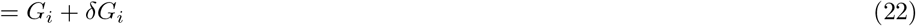

where

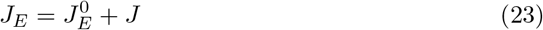

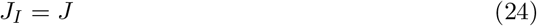

and *J >* 0, which is required for bump stability (see Eq. 68). Note that *δG* is a convolution of the approximation of 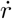 with a function composed from a combination of the DC and fundamental Fourier modes. The parameters 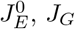 and *I*_ext_ in the term *G*_*i*_, determine the structure of the bump at steady state, whereas *J* does not have any effect on it. Feedback gain of NDF is regulated by *δG*_*i*_ via the parameter *J*. In other words, changing *J* does not affect bump structure, but does modulate the strength of NDF.

#### Multi-ring model

This model has five populations in total. For simplicity, they are assumed to have the same number of neurons *N*. Their activity is described by the following equations:

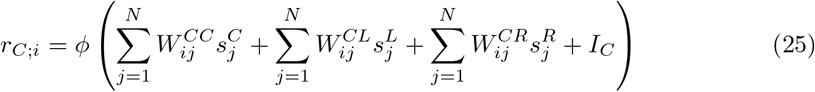

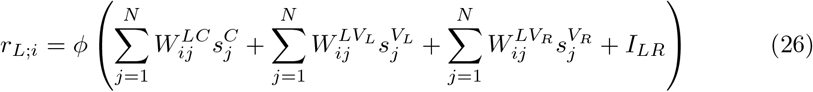

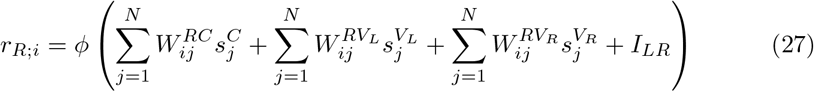

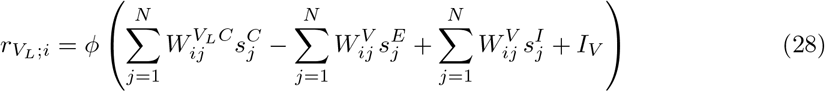

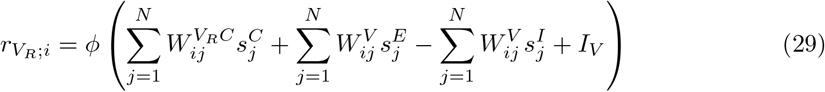

For simplicity, we assume that all synapses, other than those involved in generating the NDF signal, have the same time constant *τ*. The corresponding synaptic activations are governed by

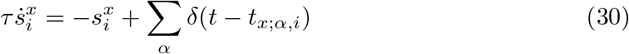

where *x* denotes the neural population, *i* denotes the neural index, and *x*∈ {*C, L, R, V*_*L*_, *V*_*R*_}. The slow and fast synaptic synapses that produce the NDF signal originate from the central population:

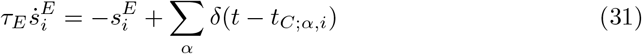

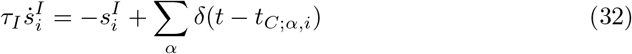

For brevity of notation, here we allow for the upper index to denote combinations of presynaptic populations and timescales (and for weights also the postsynaptic target), in difference from the convention set in the general framework.

In the multi-ring network used to compare naive with precise NDF (see Fig. 5), we use the same set of equations as above, except for the following changes: equation 25 is replaced by

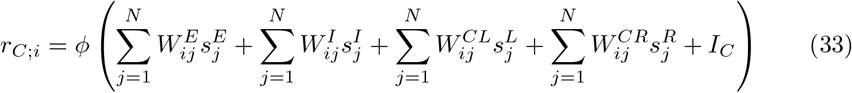

and equations 28-29 are replaced by

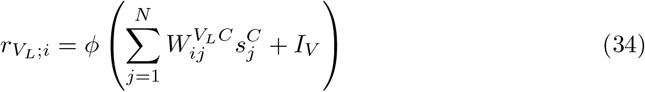

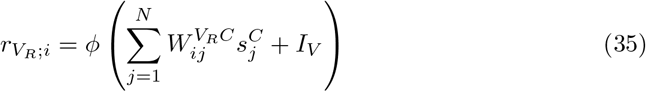

Connection matrices are given by

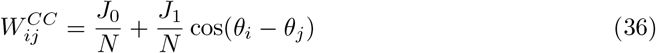

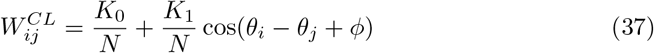

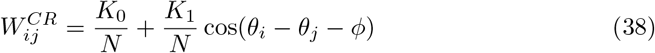

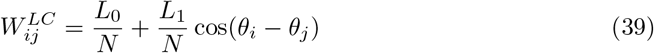

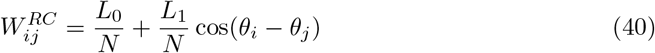

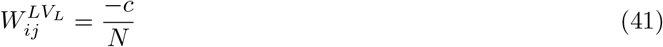

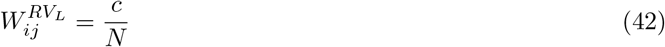

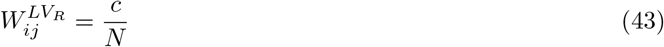

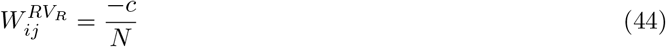

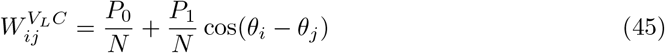

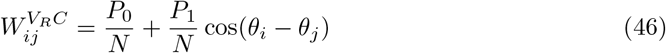

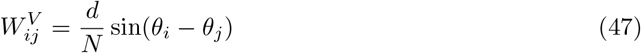

and additionally, in the multi-ring naive NDF comparison network

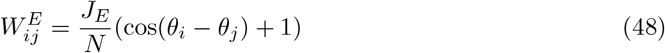

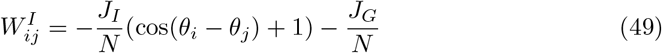

In order for both versions of the network to be equivalent when *τ*_*E*_ = *τ*_*I*_, ensuring the same diffusivity in the absence of NDF, we require that *W* ^*CC*^ = *W* ^*E*^ + *W* ^*I*^ (see 36, 48, 49), implying *J*_0_ = *J*_*E*_ − *J*_*I*_ − *J*_*G*_ and *J*_1_ = *J*_*E*_ − *J*_*I*_.

### Diffusion coefficient

Under the assumption that the attractor manifold is parametrized by the continuous variable *θ*, the diffusion coefficient is defined as

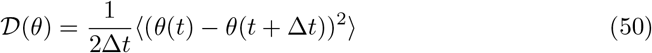

which is half of the rate of growth of the mean squared displacement of *θ*

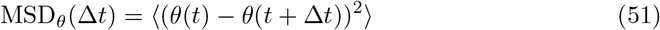

### Numerical evaluation

In the linear network, the attractor parameter was numerically evaluated as the mean firing rate

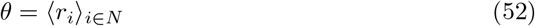

In the ring network, it was evaluated as the center of mass of firing rates:

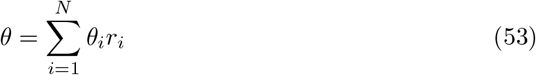

In order to avoid discontinuities, the trajectory of *θ* is unwrapped using Algorithm 1:

**Algorithm 1.**
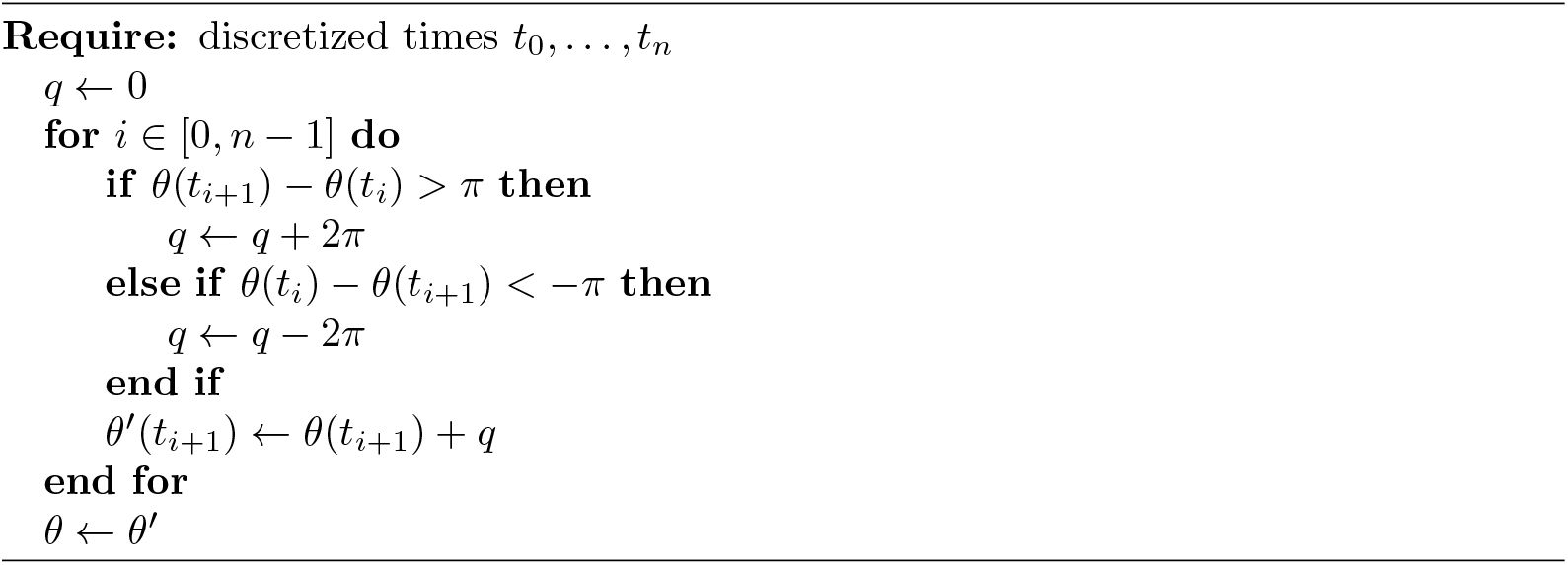
Ciruclar unwrapping.

In the multi-ring network, *θ* is evaulated from the *C*-population in the same way as it was defined in the ring network. In order to calculate a numerical estimate of the diffusion coefficient, we first obtain a large number of *θ*-trajectories (see table 1). Then, we calculate the corresponding MSD_*θ*_(Δ*t*) as per eq. 51 for an array of Δ*t*. Its slope is calculated using a linear fit. 𝒟 is calculated as half of the slope.

**Table 1.**
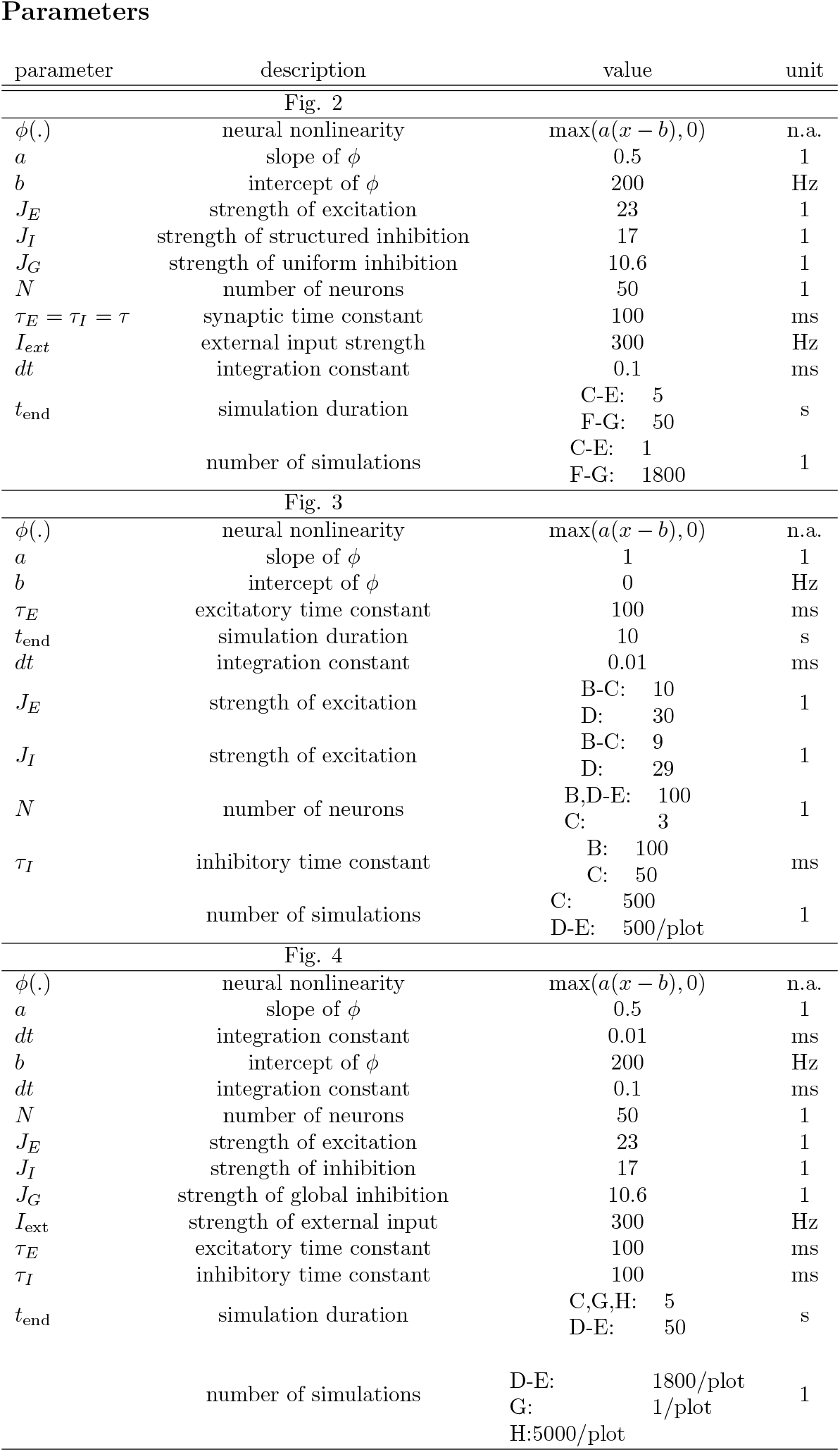
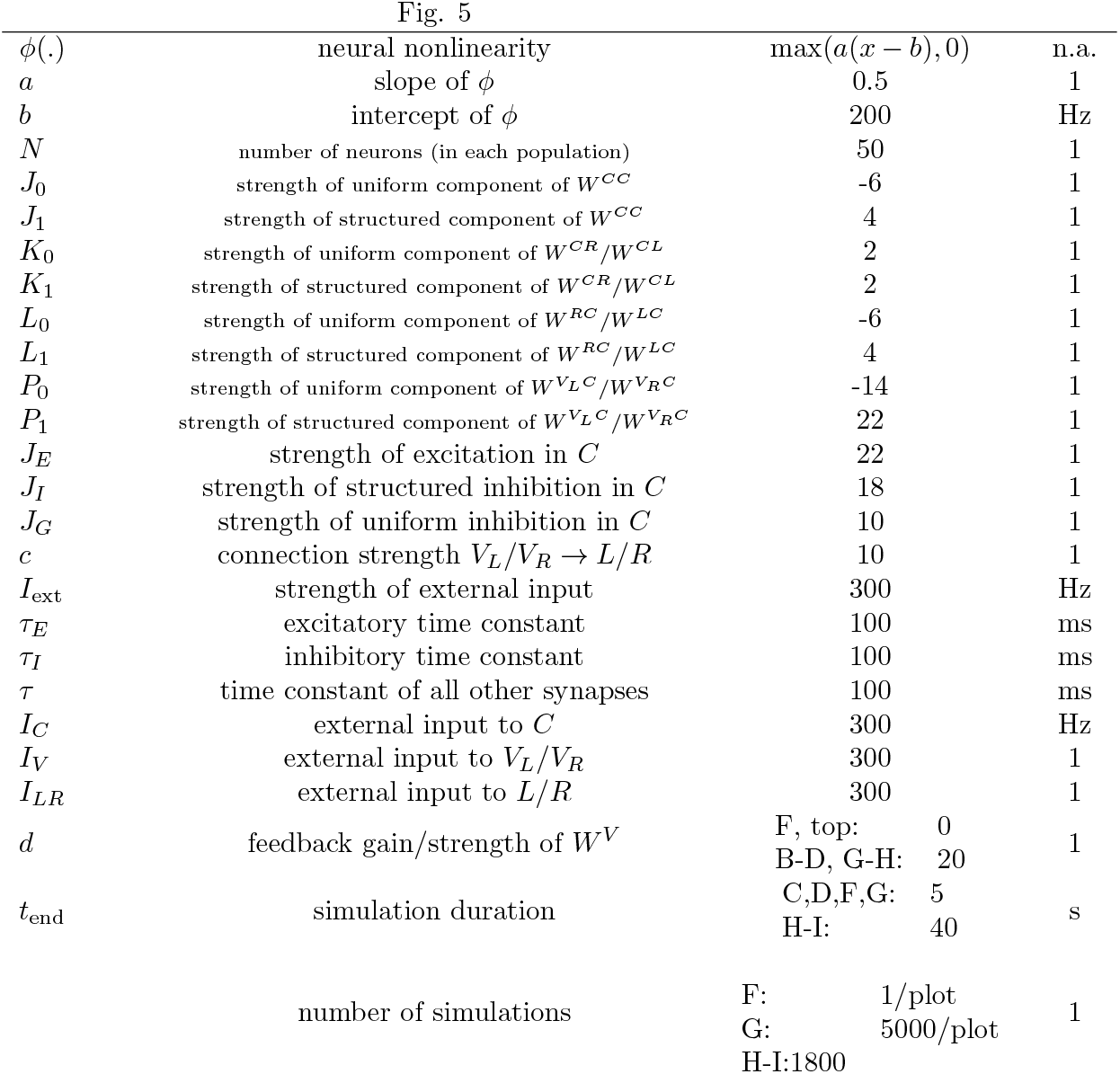
Parameters (unless specified otherwise in figure captions).

### Analytical expression for the diffusion coefficient in a noisy CAN with symmetric connectivity

For a CAN in which all connection matrices 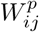 are symmetric, such as for the linear- and single population ring attractors, the diffusion coefficient is given by

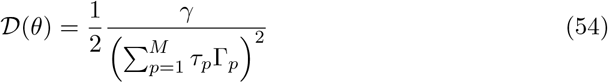

where

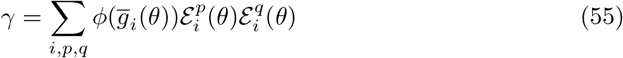

and

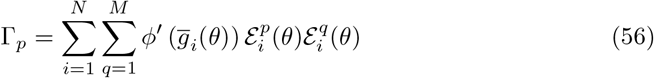

and 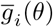 is the total synaptic input to neuron *i* (eq. 11) when the network is at the steady-state of the noise-free rate dynamics at position 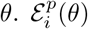 signifies the sensitivity of neuron *i* to changes in the synaptic input with timescale *p* (eq. 10) when the attractor is in position *θ* and is defined as

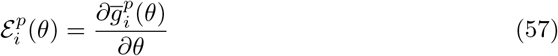

These expressions generalize results from [40] to networks with multiple synaptic timescales, and their derivation is provided in the supplementary material, D.

### Analytical expression for the diffusion coefficient in a noisy CAN with asymmetric connectivity

When the connection matrices 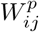 are not all symmetric, such as in the multi-ring attractor architecture, the diffusion coefficient 𝒟 can no longer be explicitly expressed analytically in terms of the steady states of the dynamics. However, it is still possible to express 𝒟 in terms of null-eigenvectors of the matrix 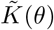,which describes the linearized dynamics of the synaptic activations 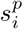, an *NM × NM* -matrix. Its left and right null-eigenvectors *v* and *u* have dimension *N×M*, but are more conveniently addressed by two indices just like 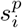.Diffusivity is then described by

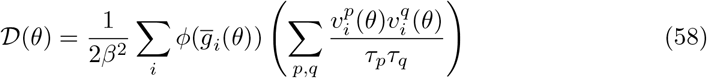

where 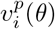 is the left eigenvector of 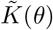,and *β* is a normalization factor that is determined by

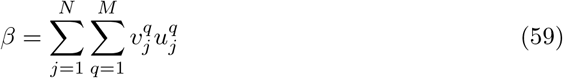

See supplementary material, D for further details.

### Negative derivative feedback and diffusion

Sufficient conditions for the reduction of diffusion via NDF in a symmetrically connected network, with distinct inhibitory and excitatory timescales, are given by the following (see Supplementary Material, D):

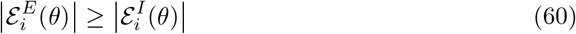

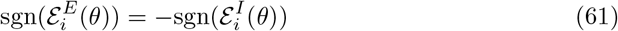

The first condition (equation 60) is that the excitatory input to neuron *i* is more sensitive to changes in the attractor parameter *θ* that the inhibitory input, and the second condition (equation 61) is that as *θ* varies, changes in the excitation and inhibition are compatible with balanced input: if the excitatory input to neuron *i* increases, so does the inhibitory input. Note that these equations imply that Γ_1_ ≤ 0 and Γ_2_ ≥ 0 in eq. 4. Under these circumstances, diffusivity will slow down as the inhibitory timescale *τ*_*I*_ is decreased relative to the excitatory one *τ*_*E*_.

It is easy to see that these conditions hold in the ring model, since the sensitivities (at *θ* = 0, without loss of generality) are given by

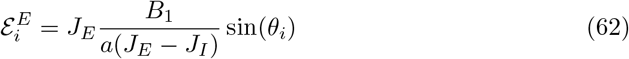

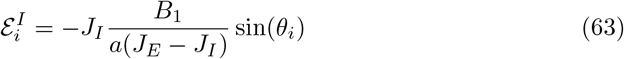

See supplementary material, Eqs. S97-S98 for the derivation of these expressions. Since

*B*_1_, *J*_*E*_, *J*_*I*_, *a >* 0, the signs of 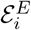 and 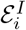 are different, and because *J*_*E*_ *> J*_*I*_ (see section on Stability Analysis), the magnitude of the former is larger than the latter.

### Linear model – analytical solution

The diffusion coefficient for the linear network is given by

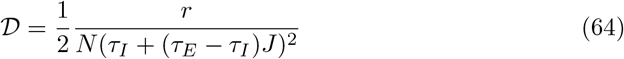

where *J* is the feedback coupling strength (and excitatory weight) and *τ*_*E*_ (*τ*_*I*_) is the excitatory (inhibitory) time constant. This result follows directly from Eq. 54.

### Ring model – analytical solution

The diffusion coefficient for the ring model can be calculated analytically and is determined by the following equation:

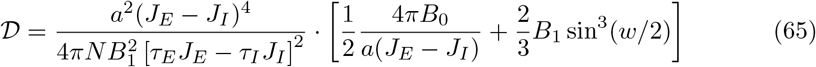

Here, *J*_*E*_ (*J*_*I*_) denotes the strength of excitation (inhibition), *a* and *b* are parameters of the ReLu function *ϕ* (Eq. 8), the excitatory (inhibitory) time constants are given by *τ*_*E*_ (*τ*_*I*_), *w* is the width of the bump, and *B*_0_ and *B*_1_ solve the equations

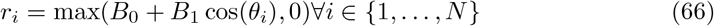

We have, without loss of generality, assumed that the attractor is in position *θ* = 0 above. See supplementary material, D for the derivation of Eq. 65 and for explicit expressions for *B*_0_, *B*_1_, and *w*.

### Ring model – stability analysis

The stability of the system’s steady states can be analyzed by linearizing the dynamics around them (see Supplementary Material, B). We find three distinct regions of stability for the parameters *J*_0_ = *J*_*E*_ − *J*_*I*_ − *J*_*G*_ and *J*_1_ = *J*_*E*_ − *J*_*I*_. If

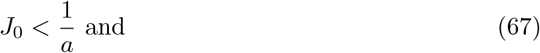

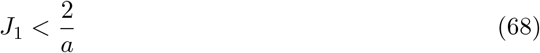

network activity is homogeneous and stable. Violation of the first condition leads to an instability in the DC-mode. Violation of the second condition leads to the emergence of a stable bump-solution in which the first Fourier component of the activity is nonzero. Thus, emergence of a stable bump state requires

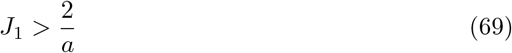

Stability of the bump state requires, in addition,

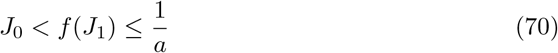

where *f* (*J*_1_) denotes the zero-crossing of the DC-mode related eigenvalue of *M* (see Fig. S1).

## Acknowledgements

This study was supported by a Synergy Grant from the European Research Council (“KILONEURONS,” grant agreement no. 951319) and the Gatsby Charitable Foundation. Y.B. is the incumbent of the William N. Skirball Chair in Neurophysics.

## Supplementary material

### A Linear model

In this section we provide details on the analysis of the linear line attractor model (Fig. 3 and Eqs. 15-17).

#### A.1 Description in terms of a two-dimensional state

It follows from Eq. 15 that all neurons in the network have the same instantaneous firing rate, which we denote by *r* (*r*_*i*_ = *r* for all *i*), and since we assume *I*_ext_ = 0,

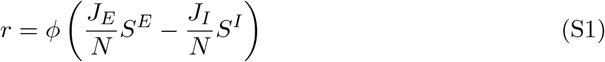

where

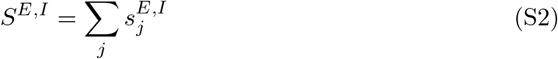

Due to Eqs. 16, 17

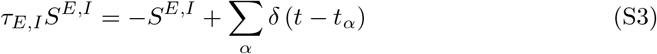

where *t*_*α*_ are the spike times of all the neurons in the network. Since each neuron produces an inhomogeneous Poisson spike train with rate *r*, the combined spike trains of all the neurons is an inhomogeneous Poisson spike train with rate *Nr*. Thus, the network is equivalent to a network with a single neuron, whose instantaneous firing rate is given by

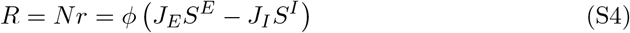

and the internal state of the system is uniquely determined by the two variables *S*^*E*^ and *S*^*I*^.

#### A.2 Condition for existence of a line attractor

In the determinstic rate model, the spike train in Eq. S3 is replaced by *r* (and in Eq. S4 by *R* = *Nr*). Thus, all neurons share the same synaptic variable 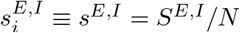.Assuming that the model operates in the linear regime of *ϕ*, the dynamics of *s*^*E*^ and *s*^*I*^ are given by

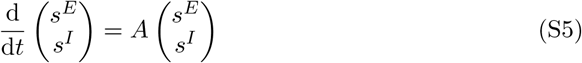

where

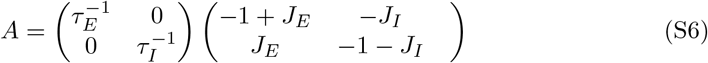

In order for the system to enter into a marginally stable regime in which the firing rate *r* can represent a memory, we need one eigenvalue of *A* to vanish, and thus

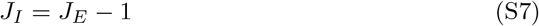

#### A.3 Negative derivative feedback

The principle through which a derivative is approximated by fast excitatory and slow inhibitory synapses can be understood as follows (see also [45]). We consider for simplicity the toy-example of a single cell with an autapse, as discussed above.

Looking at the noise-free dynamics, we have

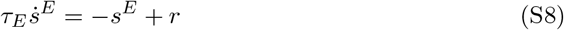

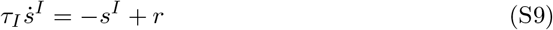

which in Fourier-space amounts to

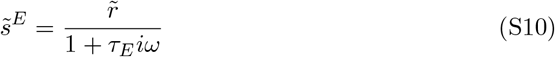

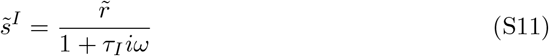

Thus,

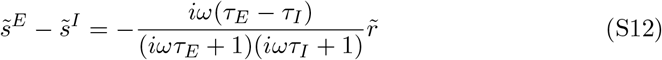

If *r* changes slowly compared to the synaptic timescales *τ*_*E*_ and *τ*_*I*_, corresponding to *ωτ*_*E*_ ≪ 1 and *ωτ*_*I*_ ≪ 1, we can conclude that

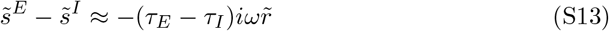

Hence,

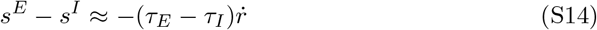

### B. Ring model

#### B.1 Steady state solution

In this section we analyze the dynamics of the ring model with two different timescales, in the absence of noise. We analytically identify the conditions for existence of a stable bump state, and the structure of the steady-state solutions. Similar analysis has previously been conducted for a ring network with a single timescale [49].

The sinusoidal structure of the weights assumed in this study (Eqs. 19-20) greatly simplifies the analysis, because the total synaptic input *g*^*i*^ takes the form of a convolution of the synaptic variables with weights that include, in Fourier space, only a constant and the fundamental frequency. Consequently, *g*_*i*_ itself is composed only of these Fourier modes, and due to the ReLu nonlinearity, *r*_*i*_ must take the form of a truncated cosine function,

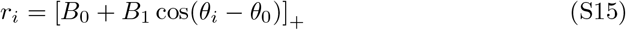

where *B*_0_, *B*_1_, and *θ*_0_ can depend on time, and are independent of time at a steady state. This parametrization can include, in principle, both a uniform state (where *B*_1_ = 0), or a bump state, in which *B*_1_ ≠ 0.

In our analysis of steady states, we shall assume, without loss of generality, that *θ*_0_ = 0 and *B*_1_ ≥ 0. Then,

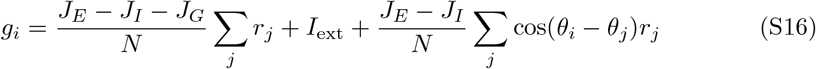

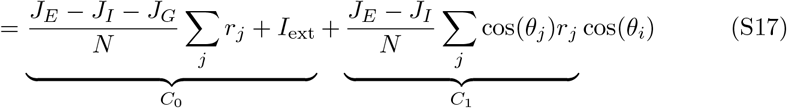

where, in the second equality, we used the symmetry of *r*_*j*_ around *θ*_*j*_ = 0. The firing rates are given by

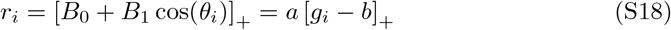

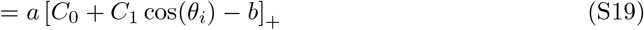

where we used Eq. 8. Thus,

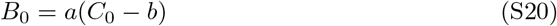

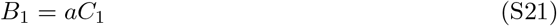

where *C*_0_ and *C*_1_ are defined below Eq. S17.

From here on, we analyze the dynamics in the limit of *N* → ∞ and replace *g*(*θ*_*i*_) by

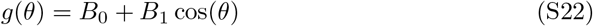

We denote by *α* the angle at which

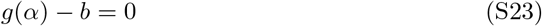

Using Eqs. S20-S23,

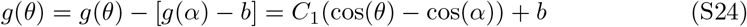

On the other hand, from S17 we see that in the large *N* limit

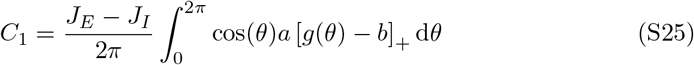

Combining Eqs. S24 and S25 we obtain

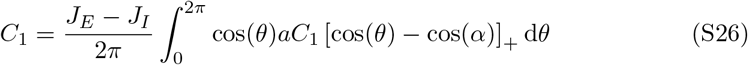

Thus,

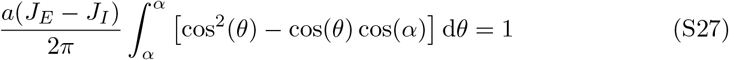

and after evaluating the integral, we get

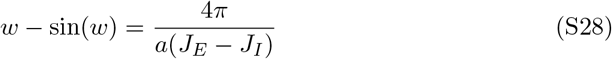

where

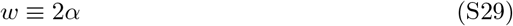

is the width of the bump. The nonlinear equation S28 can be solved numerically in order to obtain *w* as a function of *J*_*E*_, *J*_*i*_ and *a*.

Finally, we can evaluate *B*_0_ and *B*_1_ using in Eqs.S17, S20, and S21:

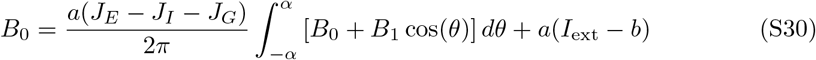

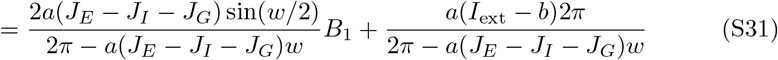

and similarly for *B*_1_:

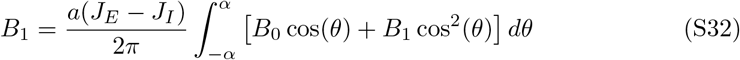

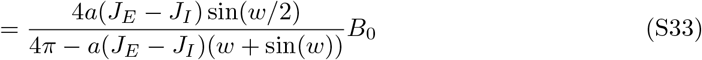

Equations S18 S28, S31, S33 together yield a complete, analytical solution of the bump shape.

#### B.2 Stability conditions

We consider the dynamics of the excitatory and inhibitory inputs *g*^*E*^ and *g*^*I*^, which are defined as

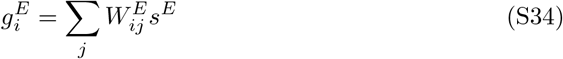

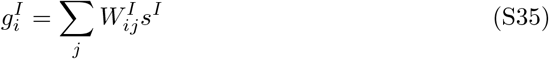

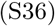

and their dynamics are

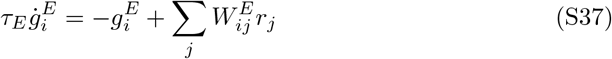

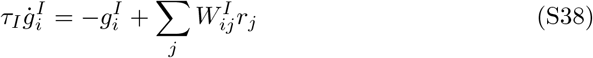

Due to the spatial structure of the synaptic weights (Eqs. 19-20), each set of inputs is composed of only three Fourier modes: the uniform mode, and the two (cosine and sine) fundamental frequency modes. Since the dynamics of the system can be uniquely re-expressed in terms of *g*^*E*^ and *g*^*I*^ [69], and each of these inputs is composed, at any given time, of three Fourier modes, the dynamical system effectively evolves within a six dimensional space.

We next linearize the system around a steady bump state, centered around *θ*_0_ = 0 without loss of generality. We determine the linearized dynamics for the Fourier-coefficients of the dynamics. For brevity, we assumed that the translational (sine) modes, which break the symmetry between clockwise and anticloskwise motion are stable, and these are excluded from the analysis presented here, resulting in a 4 × 4 dynamics matrix. However, the analysis can be readily extended to the full dynamics of the system, which then would be described by a 6 × 6 matrix. The dynamics of the system under the above assumptions are given by

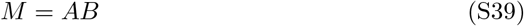

with

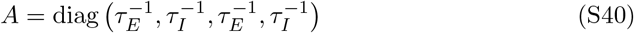

and

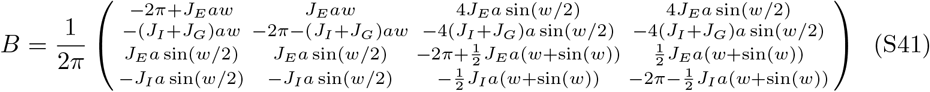

The same expressions apply also to linearization around a homogeneous stable state, which corresponds to *w* = 2*π* (whereas the case *w <* 2*π* corresponds to a localised bump state). Note that in the homogeneous case (*ω* = 0) all elements of *M* in the quadrants that couple Fourier modes with different frequencies vanish, as expected due to the rotational symmetry.

By evaluating the eigenvalues of *M* we identify three distinct regions of stability for the parameters *J*_0_ = *J*_*E*_ − *J*_*I*_ − *J*_*G*_ and *J*_1_ = *J*_*E*_ − *J*_*I*_. The homogeneous regime is stable when *J*_0_ *<* 1*/a* and *J*_1_ *<* 2*/a*. When *J*_1_ *>* 2*/a*, the homogeneous solution becomes unstable and gives way to a stable bump state so long as *J*_0_ *< f* (*J*_1_) ≤ 1*/a* (see Fig. S1), where *f* (*J*_1_) is a nonlinear function that denotes the zero-crossing of the relevant eigenvalue. Otherwise this solution is also unstable.

It is easy to see that the zero-crossings of the eigenvalues do not depend on the timescales, since

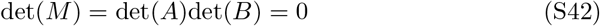

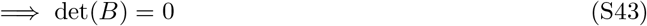

since det 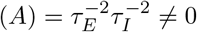 and for *τ*_*E*_ *> τ*_*I*_, the eigenvalues are real.

As can also be seen in Fig. S1, our soultion is identical to the one found in [49] and we have shown here that it is not affected by relative changes in synaptic timescales.

We note for completeness that when *τ*_*E*_ *< τ*_*I*_ the linearized dynamics around the stationary bump state can have non-real eigenvalues, and the full dynamical system can exhibit a limit cycle. In this case, the unstable modes in a linearization around the stationary bump state can involve the translational modes. Hence the analysis must be performed in the full six-dimensional subspace. We do not present this analysis here because we are concerned in the present work only with the case *τ*_*E*_ *> τ*_*i*_.

**Fig S1.**
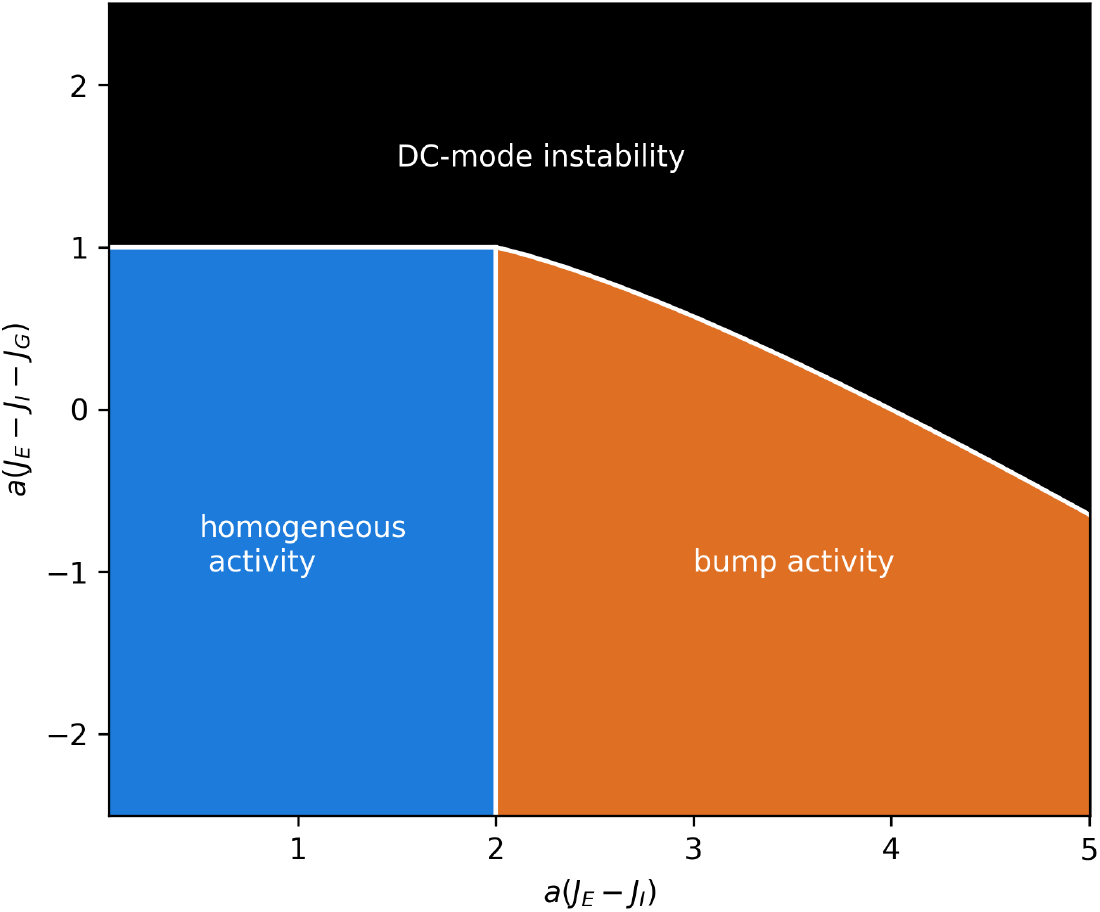
Phase diagram for the ring model.

### C Multi-ring model

#### C.1 Velocity cannot be extracted using a linear readout

To motivate the architecture of the multi-ring network, we first show that it is not possible read out the velocity of the central bump in a linear readout scheme, i.e., by projecting the derivatives of the firing rates in the central bump, onto some fixed readout vector *w*. For convenience, we move to the continuum limit and consider a general linear readout of the form

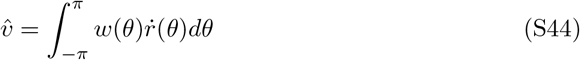

where 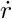 represents the derivative of the firing rate of the neuron with prefered selectivity *θ*. A fairly minimal requirement from any reasonable readout is that it should have the correct sign if an ideal bump is moving at a constant angular velocity *v*, regardless of the bump’s position on the ring. Let us denote the readout in such as a scenario as 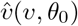,where *θ*_0_ is the current location of the bump. We then have,

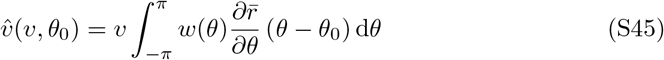

where 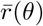 is the firing rate of the neural population in a perfect bump state (a steady state of the rate model without noise), centered at the origin. We see, consequently, that for any *v*

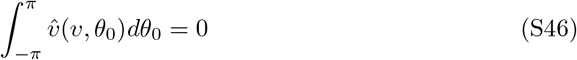

This means that if 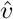 has the same sign as *v* when the bump is localized at certain positions, there will inevitably be other bump positions for which 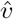 will have the wrong sign.

A related statement is that if motion on the ring is detected using a linear readout with a fixed kernel, based on *noisy* observations (e.g. from Poisson spiking neurons), there will inevitably be positions of the bump for which the Fisher information carried by 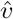 on the velocity will vanish [70].

In summary, a linear projection cannot produce a correct readout of velocity for all bump positions. However, if the bump position *θ*_0_ is known, the appropriate readout of motion is linear: the optimal readout is then obtained by projecting the synaptic activities on the left null-eigenvector of the linearized dynamics around the bump state at position *θ*_0_ [40]. The extraction of velocity is a non-linear computation because the structure of the left null-eigenvector itself varies with *θ*_0_. These insights are central to the understanding of velocity extraction in the multi-ring architecture, discussed next.

#### C.2 Extraction of velocity in the multi-ring architecture

Here we show how the layer of velocity sensing neurons extracts the bump’s velocity in an invariant manner with respect to the bump’s position *θ*_0_, based on activity in the central bump, and relays this signal into the layer of rotation cells.

We focus on the intrinsic dynamics of the central bump, and think of the velocity sensing neurons and rotation cells as comprising a separate circuit, that generates a feedback signal to counteract diffusion within the central ring. The optimal readout of velocity is then obtained by projecting the temporal derivative of synaptic activities in the central ring on the left null-eigenvector of the central bump’s intrinsic dynamics [40]. This eigenvector has the structure of a sinusoid centered at the current position of the bump. To see why this is true, note that for networks with symmetric connectivity the left null eigenvector is proportional to 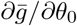,where 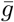 is the total synaptic input at the steady state of the noise-free dynamics in which the bump is at position *θ*_0_. The sinusoidal structure of the left-eigenvector follows from the sinusoidal structure of the recurrent weights within the central ring,

We further assume that 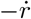 is aptly approximated by *s*^*E*^− *s*^*I*^. Thus, the desired readout of angular velocity is proportional to

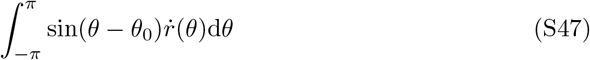

where 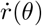 is the derivative of the firing rate of a neuron with preferred orientation *θ, θ*_0_ is the current orientation of the bump and, for simplicity, we took the continuum limit, in which a summation over discrete neuron indices is replaced by an integral over *θ* (more precisely, it is 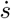 that should be projected on the sinusoidal function. For the sake of this discussion we ignore this distinction).

The firing rates of velocity sensing neurons are given by

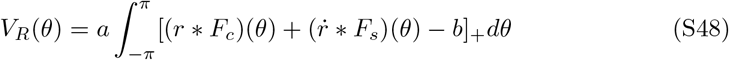

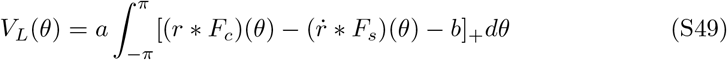

where *θ* represents the prefered angle of a neuron in the *V*_*R*_ or *V*_*L*_ population, the symbol represents the convolution operator, *F*_*c*_ is a symmetric kernel determined by the elements of the matrices 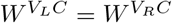,and *F*_*s*_ is an antisymmetric kernel detetmined by the elements of the matrix *W* ^*V*^ (see Eqs. 45-47). Specifically, we assume that *W* ^*V*^ is structured as a sinusoidal, Eq. 47. Taking the continuum limit,

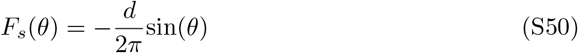

Due to the spatially uniform structure of the projections from the velocity sensing neurons to the rotation cells (Eqs. 41-44) the readout extracted by the velocity sensing neurons is

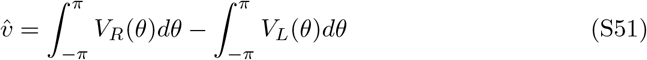

We next show that this expression implements the desired readout, Eq. S47.

We assume that the firing rates can be decomposed into those corresponding to an ideal bump at a certain position *θ*_0_, and small fluctuations arising from noise and from the local motion of the bump:

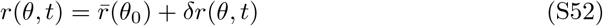

where 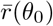 is an ideal bump state (a steady-state of the noise-free dynamics) centered around *θ*_0_ and *δr* represents the deviations from this expression. Note that our subsequent assumption that *δr* is small is applicable only if *θ*_0_ is chosen correctly. We can now linearize Eqs. S48, S49 around *r*_0_:

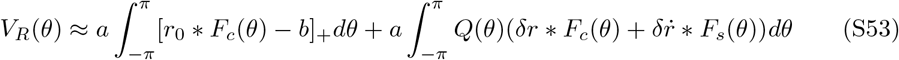

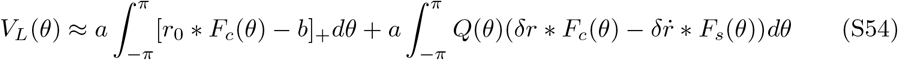

where *Q*(*θ*) = *H* [*r*_0_ ∗*F*_*c*_(*θ*) − *b*] and *H*(.) denotes the heavyside-function. It follows from Eq. S51 that the velocity estimate is given by

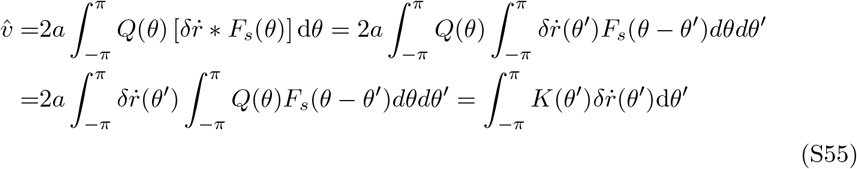

where, due to Eq. S50,

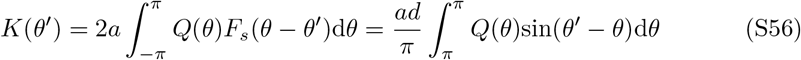

We note that *Q*(*θ*) is a positive symmetric function centered around *θ*_0_. Its circular convolution with a sinusoidal function is also a sinusoidal function, centered around *θ*_0_:

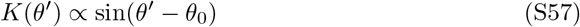

with a positive prefactor. By combining Eqs. S55 and S57 we see that 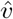 matches the desired form given by Eq. S47.

#### C.3 Alternative connectivity in the multi-ring model

In the multi-ring model, the fast and slow connections between the *C* and *V*_*L*_/*V*_*R*_-populations can be both inhibitory and excitatory. Thus, in order to decompose these connections so that they conform with Dale’s law would involve four types of connections: fast excitatory, fast inhibitory, slow excitatory and slow inhibitory. The underlying ∼ sin(*θ*_*i*_− *θ*_*j*_)-connectivty was chosen for mathematical simplicity. However, it is possible to modify this connectivty such that only two types of connections, fast inhibitory and slow excitatory are required, just like in the previous models. The modified equations for 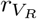 and 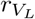 are given by:

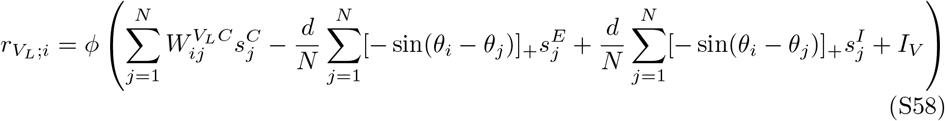

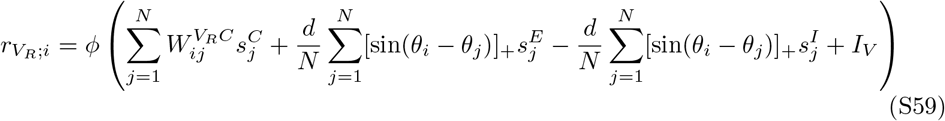

The modified connections are shown in Fig. S2A. Similarly to the multi-ring model from the main text, we achieve a reduction in diffusivity (Figs. S2B-C) while keeping the amplitude-fluctuations and amplitude relaxation time minimal (Figs. S2D-E). Note that although with this connectivity scheme the velocity is no longer read out optimally and thus *V*_*L*_ and *V*_*R*_ do react to changes in bump amplitude, they respond to such perturbations in the same way. Due to the connectivity from *V*_*L*_/*V*_*R*_ to *L*/*R*, such changes are subtracted out and do not affect the operation of the circuit, while this is not true of translational changes – those cause *V*_*L*_ and *V*_*R*_ to react in different ways, and the resulting changes are not subtracted out.

**Fig S2.**
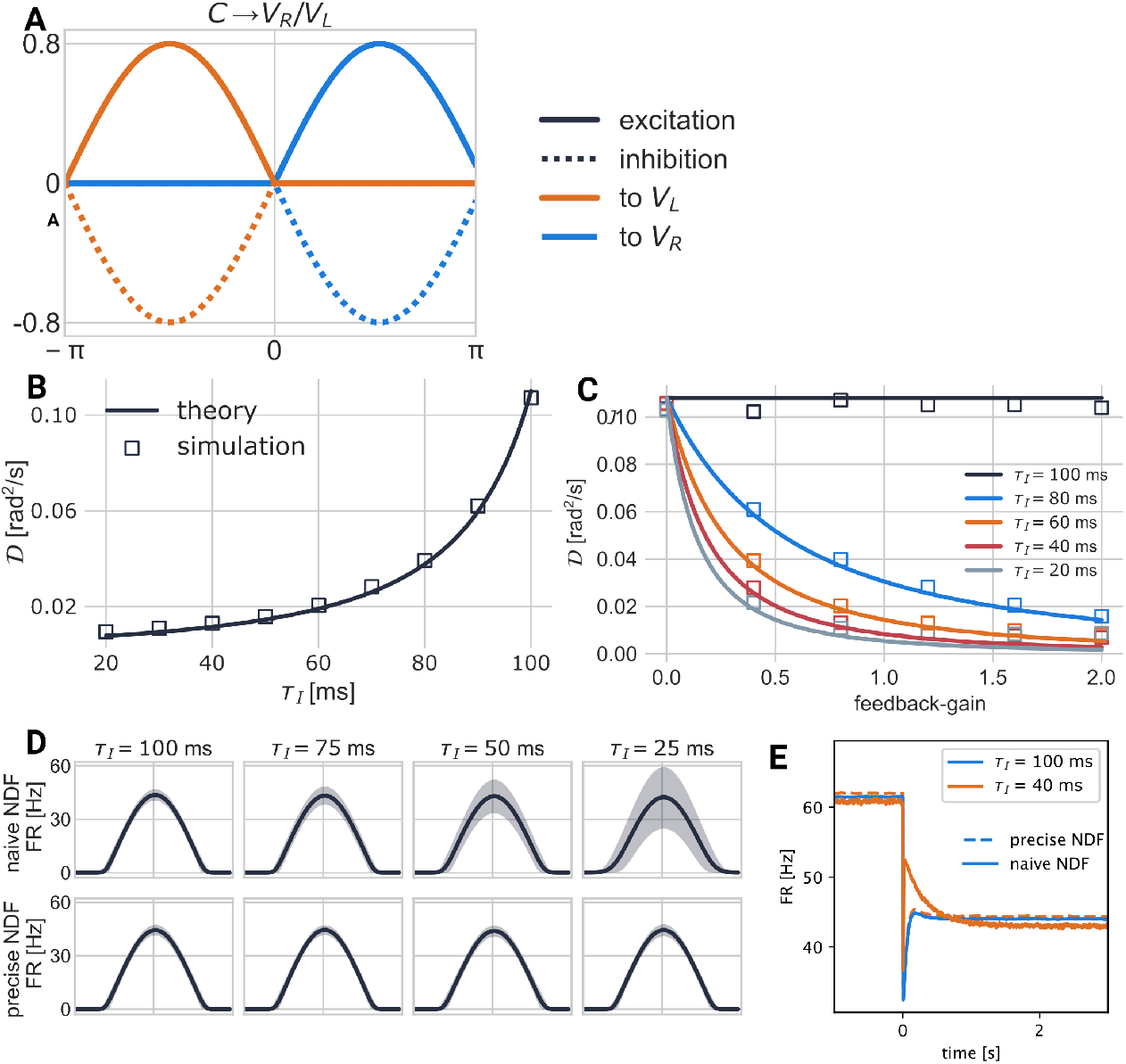
The Multi-Ring model with modified connectivity. **A**: The connections from *C* to *V*_*L*_ and *V*_*R*_. *C* projects both excitation and inhibition 90 degrees (counter-) clockwise onto *V*_*R*_ (*V*_*L*_). This connectivity fulfills Dale’s law. **B**: Diffusivity as a function of *τ*_*I*_. *τ*_*E*_ = 100 ms and feedback gain= 0.8. **C**: Diffusivity as a function of the feedback gain. **D**: Attractor tightness: variation of firing rates of the central ring. Black: mean activity. Gray shaded area represents the range of firing rates between the 0.1 and 0.9 quantiles of activity in *C*. Top row: naive NDF mechanism on central ring. Bottom row: precise NDF mechanism with modified connectivity. Left to right: decreasing *τ*_*I*_. **E**: Relaxation of bump amplitude following abrupt change in input to the central ring, at time 0, see Fig. 5G.

#### C.4 Eigenvalues elucidate relaxation dynamics

We have shown above that while naive NDF slows down relaxation into the attractor, which might be an undesired property. Another way of establishing this is to look at the eigenvalues of the corresponding (noise-free) dynamical system. We should expect that the eigenvalues which correspond to eigenvectors that pertain to changes in the amplitude of the central bump do not increase as *τ*_*I*_ becomes smaller, but instead stay the same. This would indicate that perturbations in the bump amplitude decay at the same speed, irrespective of the presence of NDF.

**Fig S3.**
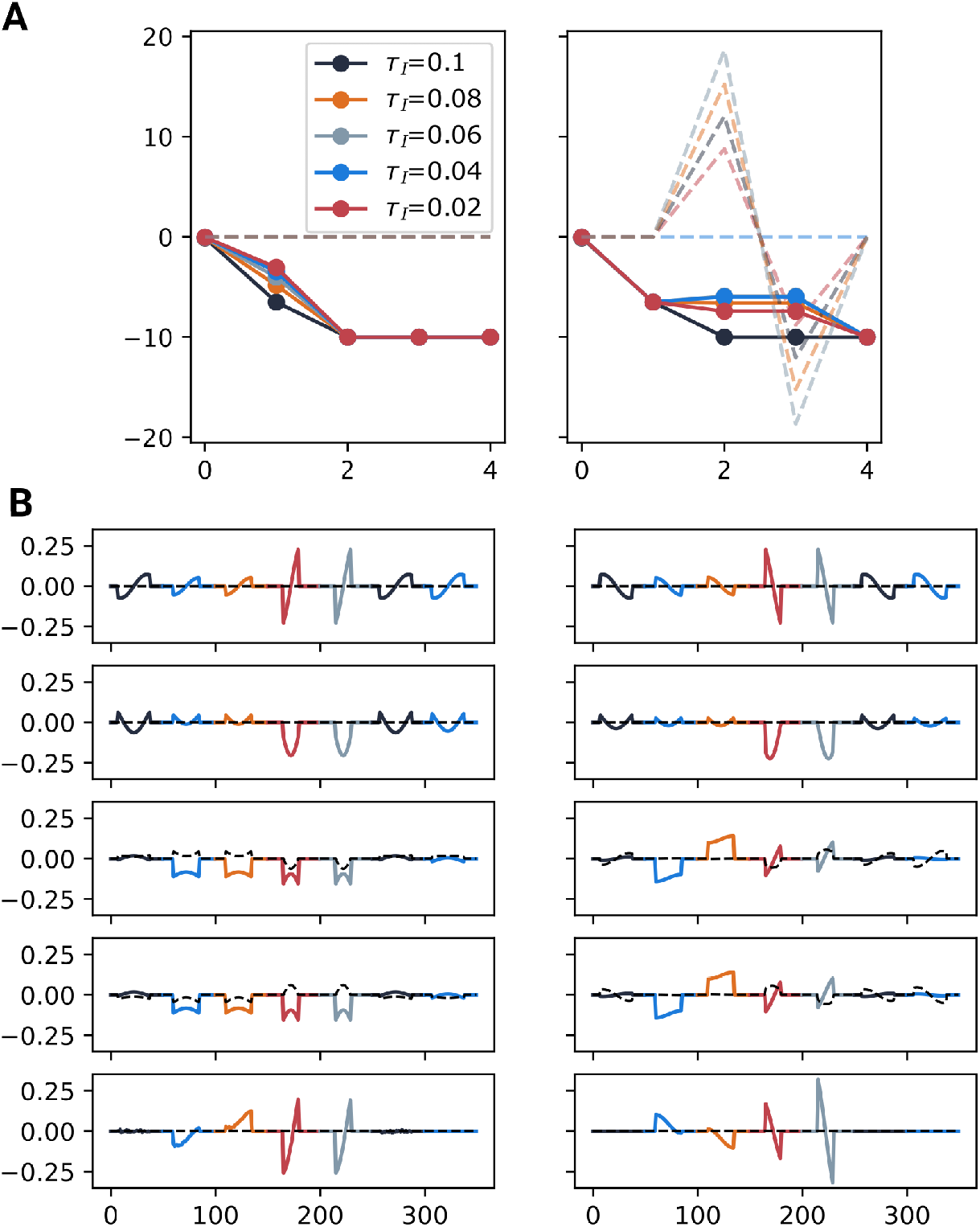
Eigenvectors and Eigenvalues in the Multi-Ring model with standard connectivity. **A**: Eigenvalues, sorted by eigenvectors (rows of B). Solid lines: real part, dashed lines: imaginary part. Left: naive NDF; second eigenvalue increases as *τ*_*I*_ decreases, indicating slowing relaxation. Note that the corresponding eigenvector (second row in left column of B) indeed corresponds to changes in bump amplitude. Right: Precise NDF; The second eigenvalue does not change as a function of *τ*_*I*_, indicating fast relaxation of amplitude perturbations (see second row of left column of B). Interestingly, the third and fourth complex-valued eigenvalues do change as a function of *τ*_*I*_, inducing small oscillations of the bump. **B**: Eigenvectors corresponding to the x-axis of A. Each color denotes a population of neurons. Black: *C*, blue: *L*, orange: *R*, red: *V*_*L*_, gray: *V*_*R*_, black, second: *C* (slow excitation), blue, second: *C* (fast inhibition). Black dashed line: imaginary part. Left: naive NDF, right: precise NDF. The first eigenvalue corresponds with the tranlational mode, and the second with changes in amplitude of the central bump. For precise NDF, the third and fourth, complex-valued eigenvectors correspond to oscillatory activity of the system.

**Fig S4.**
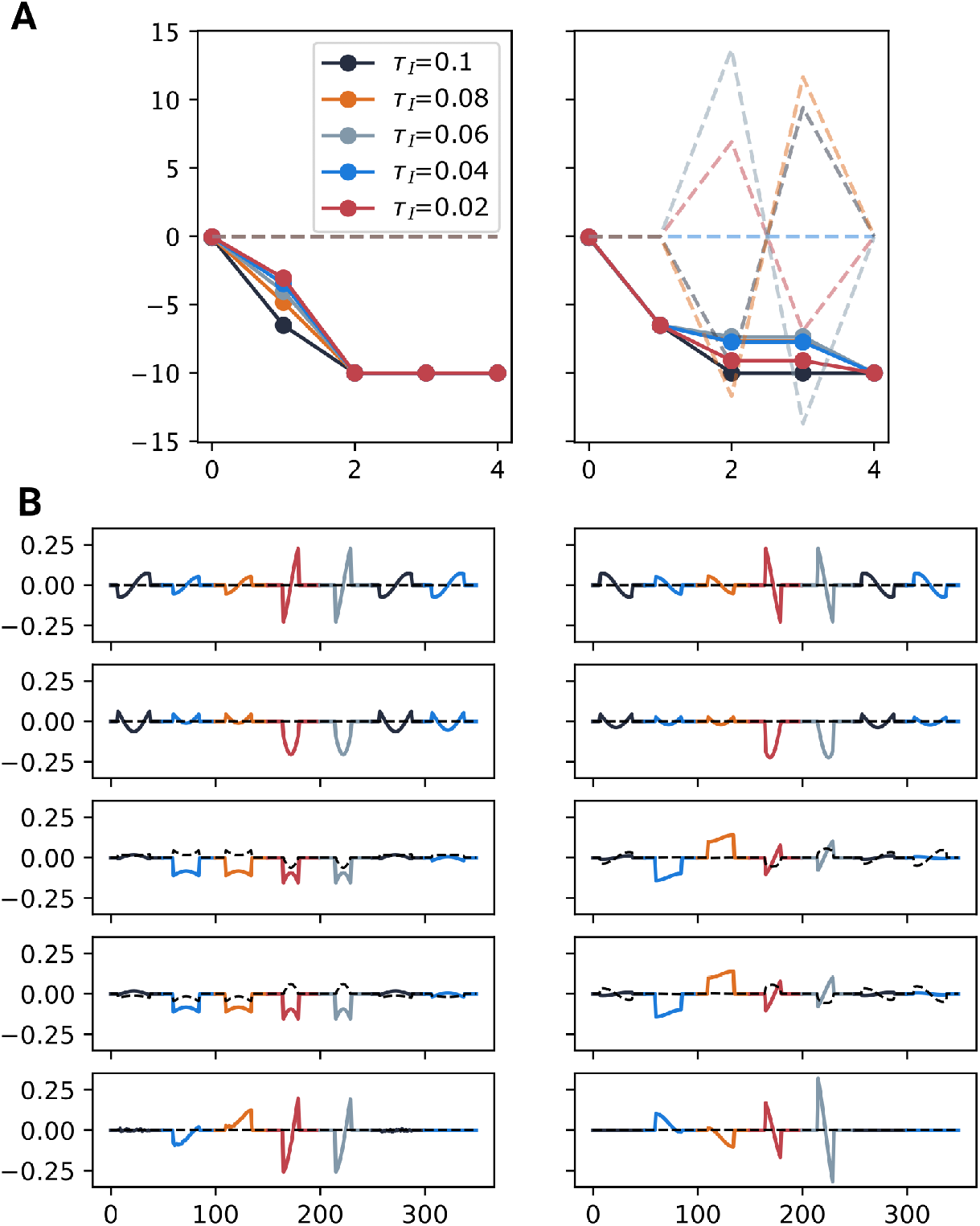
Eigenvectors and Eigenvalues in the Multi-Ring model with alternative connectivity. See S3.

Figs. S3-S4 indeed confirm this for both the standard and alternative connectivity for the multi-ring model outlined above.

**Fig S5.**
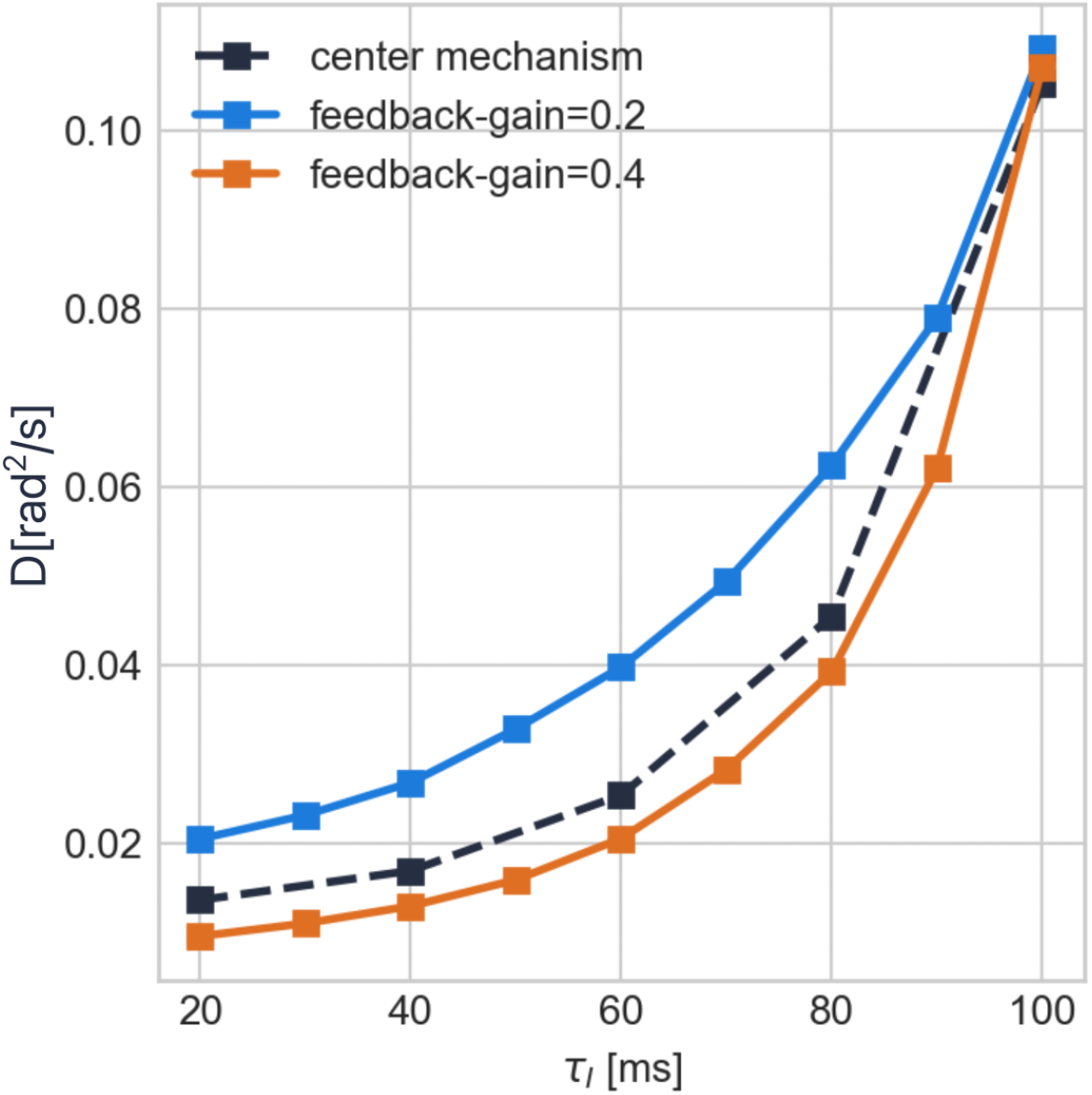
Comparison of diffusivity-curve between naive and precise NDF. For native NDF, the feedback gain was chosen such that the reduction in diffusivity lies between feedback gains of 0.2 and 0.4 for the precise NDF. Therefore, with a choice of a feedback gain of 0.4, precise NDF performs better without affecting bump variability or increasing relaxation time

### D Diffusion coefficient

#### D.1 Derivation

We calculate the diffusion coefficient for the generalized model with *N* neurons and *M* timescales. To that end, we consider the rate-approximation of the general model outlined in methods. The synaptic activation of neuron *i* with timescale *τ*_*p*_ while the attractor is in state *θ* is denoted 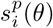.We define the state vector of the noise-free dynamics as

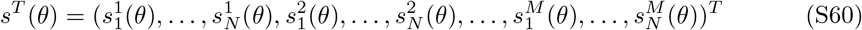

and its linearized dynamics around the steady state 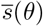 are given by

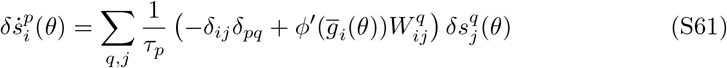

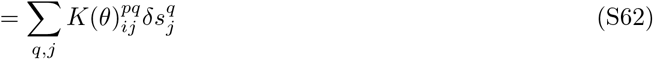

where 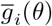 denotes the steady state of the total input *g*_*i*_(*θ*). We can write

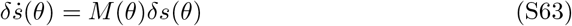

with

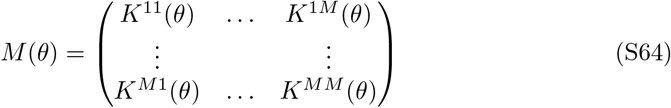

We denote the right zero-eigenvector of *M* (*θ*) *u*_0_(*θ*):

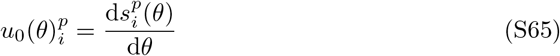

and its left zero-eigenvector *v*_0_(*θ*). We find the normalization *β* such that

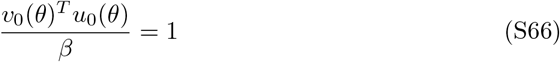

and express *δs*(*θ*) as a linear combination of its right eigenvectors:

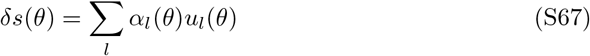

Now we reintroduce a noise term into the linearized system (see Eqs. 12-14), giving us

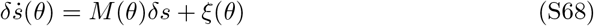

With Eqs. 12-14 and the composition of *s*(*θ*) in Eq. S60, we can see that *ξ*(*θ*) has the following structure:

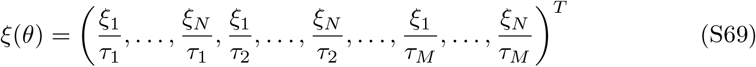

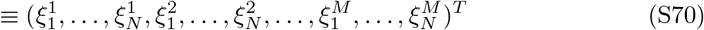

Now, by switching the indexing convention, we can express Eq. S68 as

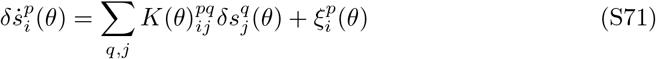

where, with Eqs. S69-S70, 14, we can see that

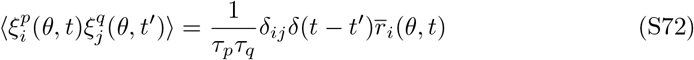

where 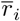. We multiply the system of equations defined by Eq. S68 and from the left by *v*_0_(*θ*)*/β* and apply Eq. S67:

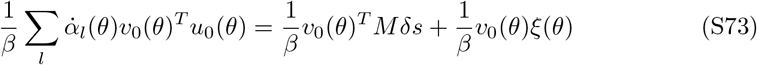

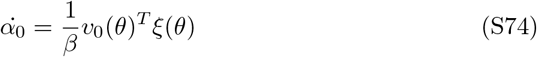

and thus

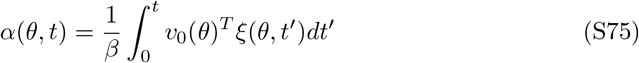

where *α*_0_ is the local approximation of the projection of the perturbation onto the attractor manifold. We can write the diffusion coefficient as

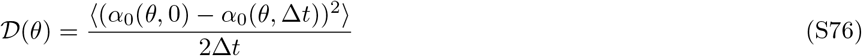

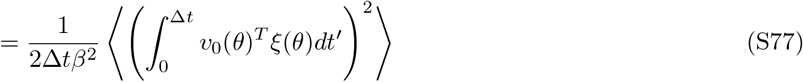

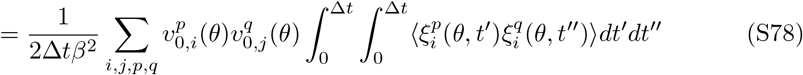

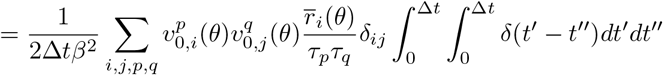

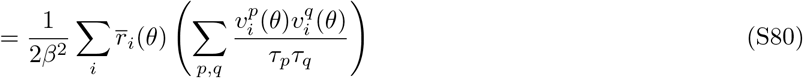

To evaluate the above expression further (at least for a special class of connectivities), we denote the elements of 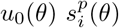.They are elements of the right zero-eigenvector of the system and therefore solve the equations

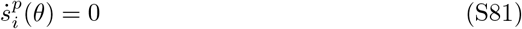

We can further evaluate this expression by noticing that

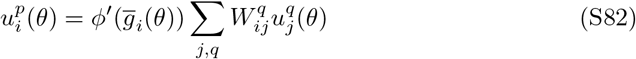

and also

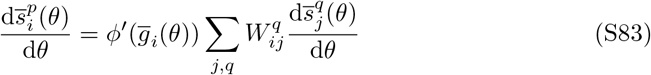

Thus we can determine that

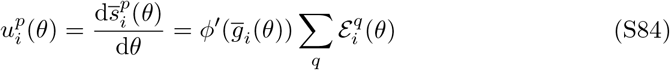

See Eq. 57 for the definition of 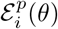.

In general, there is no anaytic expression for the left zero-eigenvector of the dynamics *v*_0_, which are determined by

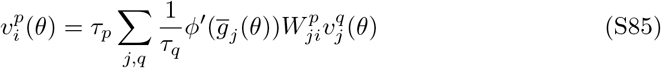

However, in the special case of a symmetric network, i.e. when *W* ^*p*^ is symmetric for all *p* ∈ [1, …, *M* ], we can obtain an analytic expression for it. The assumption of symmetry allows us to write this as

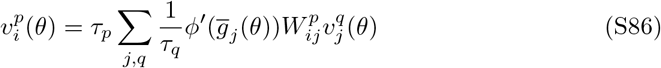

These equations are satisfied by

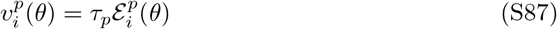

and thus,

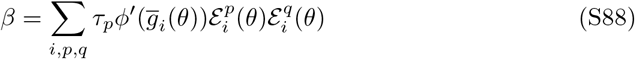

Plugging this into Eq. S80, we get

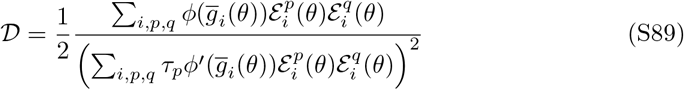

For two timescales, labelled *E* and *I*, this can be written as

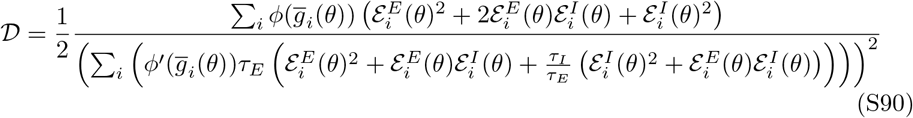

It is evident that only the denominator is affected by a difference in timescales. Furthermore, the term

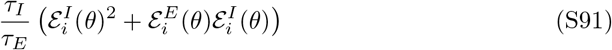

will be negative if the conditions

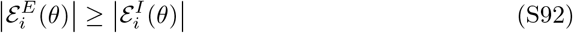

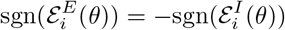

are fulfilled. If this is true for each neuron *i*, we have derived sufficient conditions to reliably reduce diffusivity, since the smaller the ratio *τ*_*I*_ */τ*_*E*_, the larger the denominator will be.

#### D.2 Analytical derivation of the diffusivity in the ring Model

Eq. S89 applies to the ring model since its connectivity is symmetric. We can explicitly calculate the diffusion coefficient by exploiting the fact that we have an analytic solution for its steady state (see above). Building on Eq. S18, we note that

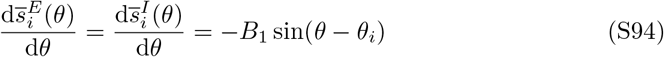

if *r*_*i*_(*θ*) *>* 0, and 0 else which, under the assumption *θ* = 0 (without loss of generality, since diffusion is uniform throughout the ring and no angle is special) becomes

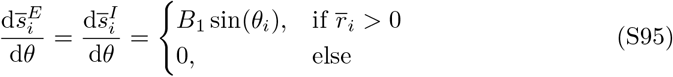

We can calculate the sensitivities as

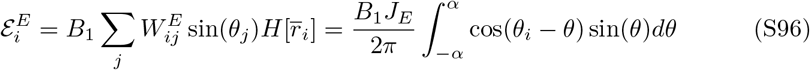

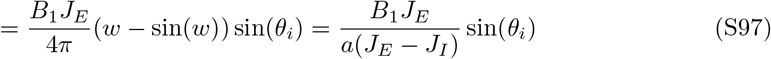

where *H*[*x*] is the Heavyside-function of *x*, ± *α* are the angles at which the zero-centered firing rates vanish and *w* = 2*α* is the bump width. We also made use of Eq. S28. Similarly, we get

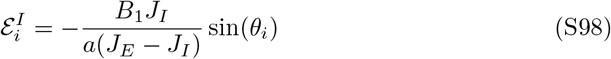

For better legibility, we write Eq. S89 as

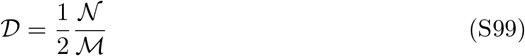

and for the ring, we get

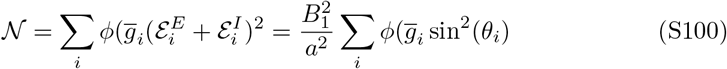

and

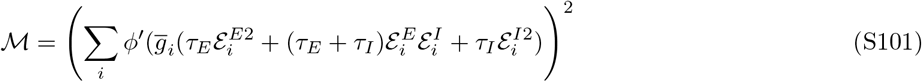

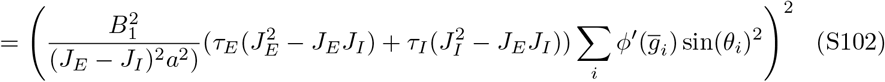

and

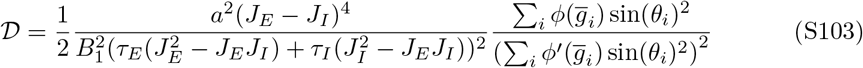

We can calculate the second factor as

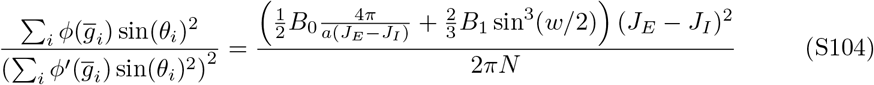

So in total

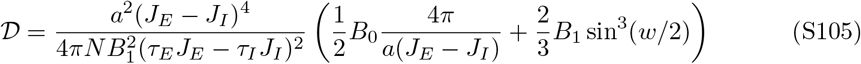

We see that diffusivity scales with *N* ^−1^, as shown in [40].

Furthermore, the dependency on timescales is clear: the smaller *τ*_*I*_ becomes, the larger the denominator and the more diffusivity gets reduced.

In order to investigate the dependence on timescales and feedback gain, we note that in order to modulate feedback gain without changing the bump shape, both *J*_*E*_ − *J*_*I*_ and *J*_*E*_ − *J*_*I*_ − *J*_*G*_ must be kept constant (see Eqs. S18, S28, S31, S33). We adapt the following parametrization of feedback gain that accomplishes this:

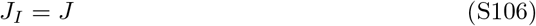

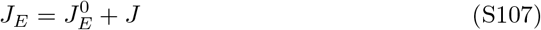

where large *J* means strong feedback gain, and 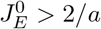,in accordance with he stability conditions. We then note that the only term that changes as a function of *J* is (*τ*_*E*_*J*_*E*_ − *τ*_*I*_ *J*_*I*_)^2^ in the denominator of 𝒟. We rewrite it as

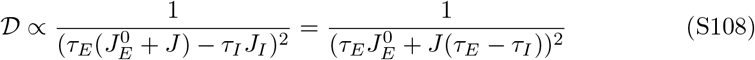

Now it is easy to see that decreasing *τ*_*I*_ relative to *τ*_*E*_ will grow the denominator, resulting in weaker diffusivity. Also, as long as *τ*_*I*_ *< τ*_*E*_, increasing *J* will lead to a decrease in diffusivity.

